# Kinetic landscape of single virus-like particles highlights the efficacy of SARS-Cov-2 internalization

**DOI:** 10.1101/2024.06.10.598174

**Authors:** Aleksandar Atemin, Aneliya Ivanova, Wiley Peppel, Rumen Stamatov, Rodrigo Gallegos, Haley Durden, Sonya Uzunova, Michael D. Vershinin, Saveez Saffarian, Stoyno S. Stoynov

## Abstract

The efficiency of virus internalization into target cells is a major determinant of infectivity. SARS-CoV-2 internalization occurs via S-protein-mediated cell binding followed either by direct fusion with the plasma membrane or endocytosis and subsequent fusion with the endosomal membrane. Despite the crucial role of virus internalization, the precise kinetics of the processes involved remains elusive. We developed a pipeline, which combines live-cell microscopy and advanced image analysis, for measuring the rates of multiple internalization-associated molecular events of single SARS-CoV-2-virus-like particles (VLPs), including endosome ingression, pH change, and nucleocapsid release. Our live-cell imaging experiments demonstrate that only a few minutes after binding to the plasma membrane, VLPs ingress into Rab5-negative endosomes via Dynamin-dependent scission. Less than two minutes later, the pH of VLPs drops below 5 followed by an increase in VLP speed, yet these two events are not interrelated. Nucleocapsid release from the VLPs occurs with similar kinetics to the pH drop, suggesting that VLP fusion occurs during endosome acidification. Neither Omicron mutations nor abrogation of the S protein polybasic cleavage site altered the rate of VLP internalization events, indicating that they do not affect these processes. Finally, we observe that VLP internalization occurs two to three times faster in VeroE6 than in A549 cells, which may contribute to the greater susceptibility of the former cell line to SARS-CoV-2 infection. Taken together, our precise measurements of the kinetics of VLP internalization-associated processes shed light on their contribution to the effectiveness of SARS-CoV-2 propagation in cells. Time-lapse videos of the studied internalization events can be accessed in the dedicated [COVIDynamics database](https://covidynamics.imb.bas.bg/).

## Introduction

COVID19 caused by the SARS-CoV-2 virus led to a major disruption of everyday life, demonstrating how zoonotic events can occur unexpectedly and have a catastrophic impact on the general population^1,2^. At present, more than 774 million cases have been reported, of which more than 7 million are reported to be fatal (WHO). Part of the β-coronavirus family^3,4^, SARS-CoV-2 is a positive-sense single-stranded RNA virus harboring 14 ORFs that encode 27 proteins^5^. These include structural proteins, namely, the S (spike), M (membrane), N (nucleocapsid), and E (envelope), which play major roles in viral survival, propagation, infectivity, and virulence^6,7^. Different systems have been developed to recreate the SARS-CoV-2 infection process in low-biosafety lab conditions. A prominent approach is the use of virus-like particles (VLPs)^8–13^ which self-assemble in cells after expression of the SARS-CoV-2 structural proteins. VLPs faithfully recapitulate the step of viral entry, while unable to replicate, which renders them non-infectious^8^. SARS-CoV-2 VLP formation is driven primarily by the M protein, while the E protein plays a potentiating role ^10,12^. While these two proteins are sufficient to form a particle, the S protein is required for entry into host cells. Interestingly, the addition of S to SARS-CoV-2 VLPs leads to a decrease in the amount of M and E. The optimal proportion of structural proteins M, N, E, and S for the production of stable SARS-CoV-2 VLPs is 3:12:2:5^10,12^. Further, addition of a specific cis-acting RNA element derived from SARS-CoV-2 to the VLPs increased packaging efficiency^8^.

Two mechanisms, namely, membrane fusion and endocytosis, have been demonstrated to play a role in SARS-CoV-2 entry into cells^14^. SARS-CoV-2 binds to target cell membranes through the interaction of S with cell membrane receptors (SR-B1, AXL, KIM1/TIM1, CD147, Neuropilin-1,2, DC-SIGN, L-SIGN, and others), the most prominent of which is angiotensin-converting enzyme 2 (ACE2)^14–25^. A considerable number of receptors and receptor co-factors have been proven as targets for the SARS-CoV-2 virus, accounting for its wide tissue and cell tropism. The S protein-ACE2 complex is recognized by the membrane-bound transmembrane serine protease 2 (TRMPRSS2), which cleaves the S protein at the S2′ site, leading to a dramatic conformational change that allows fusion between the virus and host cell membranes^14^. This results in the release of viral genetic information into the cell, setting the stage for viral replication^26^. If the S protein-receptor complex is not engaged by this protease due to a low membrane concentration or absence of the latter, ACE2-bound SARS-CoV-2 is internalised via clathrin-mediated endocytosis^27^. The pH of the virus-containing endosomes then drops, leading to activation of the cathepsin L protease^28^. Cathepsin L-mediated cleavage of SARS-CoV-2 at the S2′ site enables fusion of the viral and endosome membranes as well as nucleocapsid release into the host cell^29^.

Recent real-time imaging studies provided valuable insight into the process of SARS-CoV-2 internalization. Tracking of single vesicular stomatitis virus (VSV) chimeras containing the SARS-CoV-2 S protein revealed that SARS-CoV-2 entry requires an acidic environment^30,31^. A detailed understanding of viral entry, however, requires a comparison of kinetic parameters of VLP internalization to those of acidification and endosomal ingress, which requires VLPs possessing all four SARS-CoV-2 structural proteins.

Herein, we employed live-cell imaging and a dedicated image analysis pipeline (**S**ingle-**Par**ticle **T**racking **A**nalysis in **C**ells **U**sing **S**oftware **S**olutions, SPARTACUSS) to precisely follow and quantify the timing of VLP internalization, VLP-containing endosome ingression, acidification, active microtubular transport, and nucleocapsid release. Our temporal characterization reveals the sequence and interdependence of the above-described processes. Rapid VLP acidification which coincides with Dynamin-mediated endosome scission and nucleocapsid release occurs 4 and 12 minutes after plasma membrane binding in VeroE6 and A549 cells, respectively, quickly followed by the initiation of active microtubule-dependent VLP motion. Our results suggest that VLP fusion (nucleocapsid release) occurs in parallel to or shortly after endosome formation. Surprisingly, the VLPs do not colocalize with early endosomes during VLP internalization. The more rapid internalization observed in VeroE6 cells may contribute to the infectivity of SARS-CoV-2 observed in these cells relative to A549. Further, neither Omicron nor del-1 mutations influence the kinetics of SARS-CoV-2 VLP internalization.

## Results

### Visualization of VLP internalization

To study the kinetics of SARS-CoV-2 entry into cells, we employed VLPs derived from HEK293 cells overexpressing SARS-CoV-2 structural proteins M, E, and S of the Wuhan variant^11^. These VLPs successfully recapitulate SARS-CoV-2 internalization^10,12^, binding to the host cell surface via S protein, whereafter they are internalized, as shown by a TEM micrograph of thin-sliced MLE-12 cells in the process of endocytosing a SARS-CoV-2 VLP^Wu^ (Extended Data Fig.1). To visualize internalization, we used VLPs containing M-mCherry (as well as unlabeled M)^12^. Hereafter, VLP^Wu^:M^Ch^ would be short for VLP^Wuhan^:(E, S, M&M-mCherry). We treated U2OS cells which overexpress NeonGreen-tagged ACE2, non-labelled ACE2 and TRMPRSS2 with these VLPs and visualized the movement of particles in 3D at 30-s intervals via multi-point spinning-disk live-cell microscopy. We observed adherence of the VLPs to the cell membrane, often on the filopodia, as previously suggested^32^ (Fig. 1A, Video S1-S3). We manually tracked each particle after membrane binding to acquire its speed and position in 3D (see Materials and Methods). Interpreting these multidimensional data requires clear and intuitive visualization, as well as precise measurement. To this end, we coupled the tracking procedure to a post-processing pipeline for extracting and visualising the multidimensional tracking results for each single VLP. We named this procedure SPARTACUSS (Single PARTicle Tracking Analysis in Cells Using Software Solutions).

**Fig. 1.**
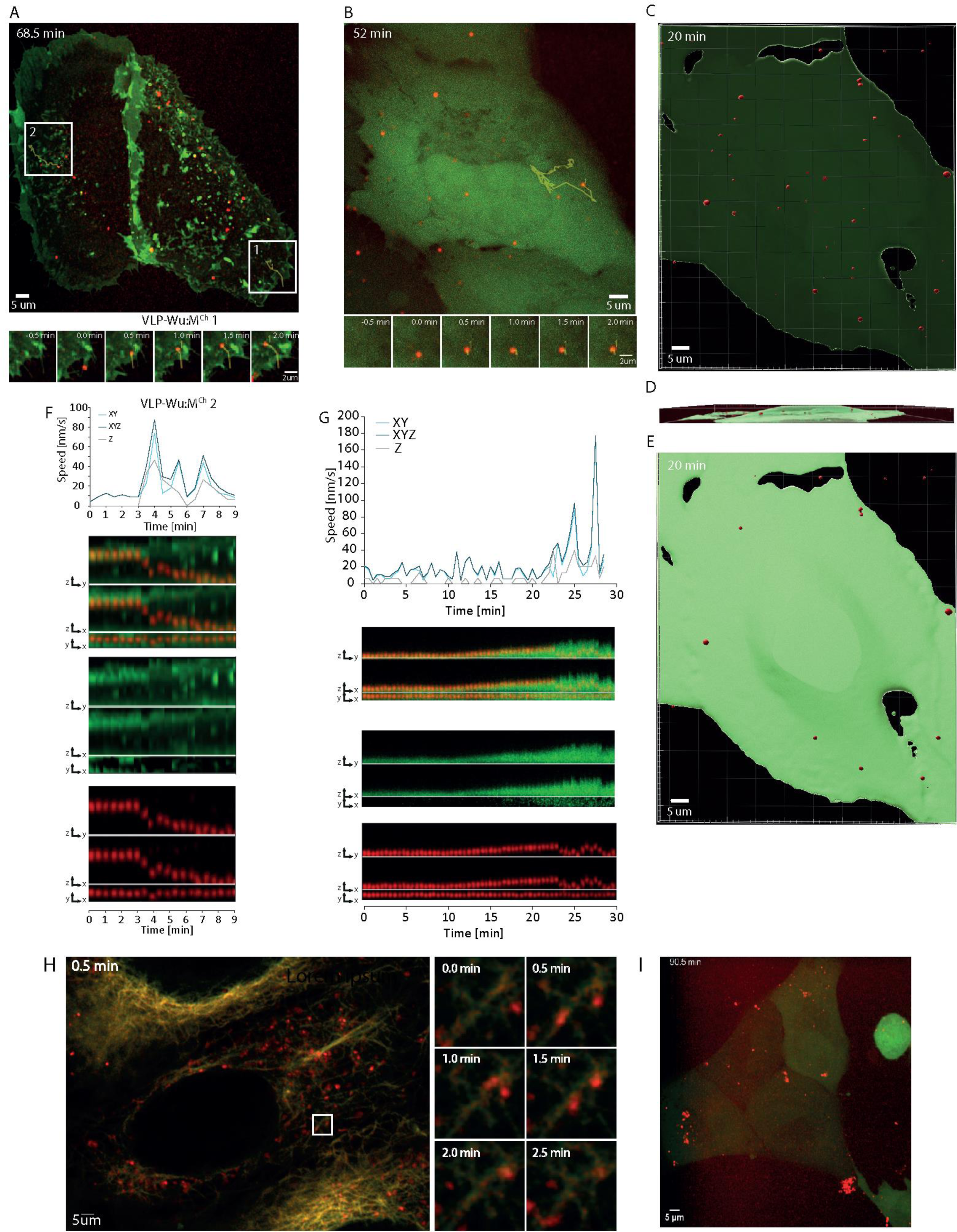
Visualization of SARS-CoV-2 VLPs binding and internalization into host cells. A. Binding and internalization of SARS-CoV-2 VLP^Wu^:M^Ch^ into U2OS cells overexpressing ACE2-Neon green and TMPRSS2. The montage shows VLPs binding to filopodia first and then migrating into the cell body. B. U2OS cells expressing mNeonGreen (to visualize cell volume) treated with VLP^Wu^:M^Ch^. C. The positions of VLP^Wu^:M^Ch^ in the transparent 3D volume of a U2OS cell (viewed from the top). D. The cell from (C) shown from the side. E. Same as (C) except VLP^Wu^:M^Ch^ are shown in a volumized U2OS cell to highlight internalized VLPs (viewed from the top) F. Graph showing the speed of the VLP (2) from (A), each point in the graph represents the speed of the particle over time extracted from two consecutive images in 1D - vertically (Z), 2D (XY), and 3D (XYZ). The kymograph represents the change in intensity of the particle and its position in Z as described in detail in Extended Data Fig.1. The time scales of the graph and kymographs are aligned. G. Same as (F), but for the particle in (B). H. Cells treated with Tubulin 610 conjugated to cabazitaxel, showing the transport of VLP^Wu^:M^Ch^ via microtubules. I. SARS-CoV-2 VLPs-WT M treated with a Recombinant Anti-SARS-CoV-2 S1 antibody. The VLPs aggregate and are unable to internalize into cells.

For the visualisation of a single particle via SPARTACUSS, we first crop the immediate space around the particle’s x and y coordinates while keeping the entire z-stack, additionally we used Gaussian blur to make the VLPs more distinct (Extended Data Fig. 2A). Next, we perform maximum intensity projections as follows: maximum intensity of x- to show the particle’s position in z and y; maximum intensity of y – to visualise the particle’s position in z and x; maximum intensity of z – to depict the particle’s position in x and y. We repeat this for each time point and combine the crops in a one-block kymograph (Extended Data Fig. 2B, C). Finally, we create such a block for the 488 channel (Green), 591 mCherry (Red) channel, and a merged block for both channels (Extended Data Fig. 2C). SPARTACUSS allows us to measure and plot the speed in 3D (x,y,z), as well as the changes in the intensity of the labelled structural proteins in individual VLPs (Extended Data Fig. 2C)

Using the SPARTACUSS workflow, we observed that, after binding, the VLPs initially moved slower (10-20 nm/s) whereafter their speed increased. This speed increase frequently coincided with downward motion in z, suggestive of particle internalization (Fig 1-A, B, F, G, Video S1-S7).

To confirm if this downward motion indeed marks VLP entry into cells, we expressed mNeonGreen, which freely diffuses throughout the whole cell volume, using it to reconstruct the cells in 3D (Video S4-S6). To see both the membrane-bound and internalized VLPs, we use transparent visualization of the mNeonGreen cell volume (Fig. 1-C, Video S7). In order to distinguish between the two VLP populations, we make the volume opaque (Fig. 1-E, Video S7), which renders most VLPs not visible once inside the cell. Internalized VLPs can be subsequently visualized using a side view of the transparent cell volume image (Fig. 1-D, Video S7). Visualization of a single VLP confirms that its speed increase in 3D coincides with cell surface penetration, as observed with the ACE2 tagging discussed above (Fig. 1-B, G). These results clearly demonstrate the capability of SPARTACUSS to capture the exact moment of VLP entry into cells and thus measure its 3D dynamics in real time.

To understand if active transport along microtubules underpins the increased movement of particles once inside the cell, we pre-incubated cells with VLP^Wu^ :M^Ch^, treated them with Aberrior LIVE 610 tubulin dye (cabazitaxel conjugated with LIVE 610 dye) for 10 min and immediately proceeded with live-cell imaging. We observed continuous co-localization between VLPs and microtubules (Fig. 1-H), indicating that fast particle movement occurs along the microtubule network (Video S8).

Next, we sought to assess how anti-SARS-CoV-2 S protein antibodies neutralize the SARS-CoV-2 virus. To this end, we incubated cells with VLP^Wu^:M^Ch^ pre-treated with an Anti-SARS-CoV-2 Spike S1 (CR3022 clone) antibody. Antibody pre-incubation precipitated most of the VLPs, preventing cell entry by effectively reducing their free concentration (Fig. 1I and Video S9)

### Dynamin-dependent VLP entry

Endocytosis has been suggested to play a role in VLP entry^33,34^. To determine the duration of time between VLP binding and ingress, we expressed GFP-tagged Dynamin in Vero E6 cells which are highly susceptible to SARS-CoV-2 infection ^35^. Dynamin binds to the invaginated clathrin-coated vesicles for 10 to 15 s and is responsible for vesicle scission during endocytosis^36–38^. The transient dynamin foci formed during this process are a standard marker of endosome formation^36,39–41^. Time-lapse imaging of Dynamin-1-GFP-expressing cells treated with VLP^Wu^:M^Ch^ allowed us to visualize VLP movement across the cells as well as the “blinking” of short-lived Dynamin foci, indicative for endosome ingress (Video S10, S11, S12). 49% of the VLPs co-localized with Dynamin foci during or immediately after the above-described speed increase and downward movement (Fig. 2A, B). This transient colocalization strongly suggests that endocytosis is involved in VLP internalization.

**Fig. 2.**
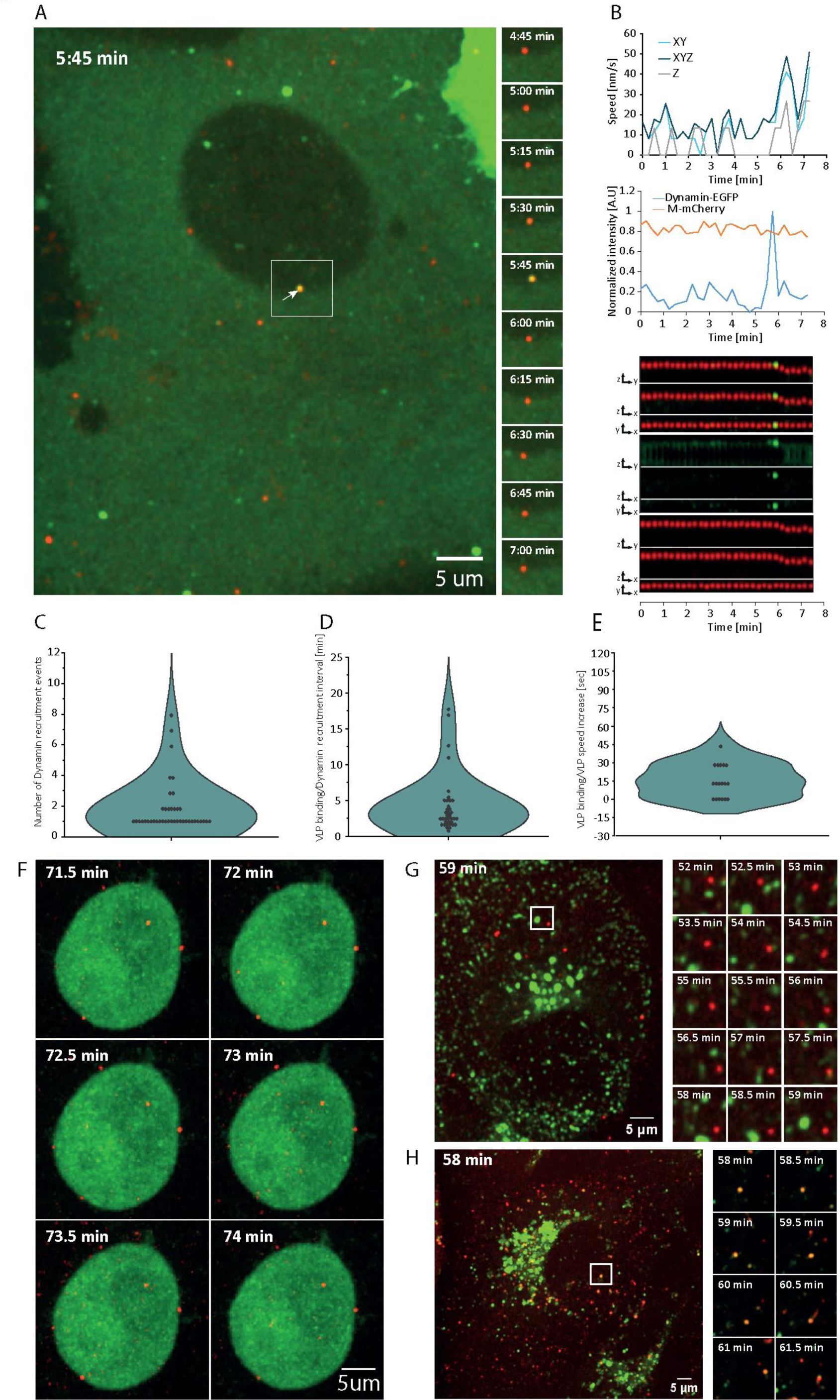
Dynamics of dynamin recruitment to the site of SARS-CoV-2 VLP^Wu^:M^Ch^ binding in Vero E6 cells. A. A representative image of SARS-CoV-2 VLP^Wu^:M^Ch^ added to Vero E6 cells and dynamin recruitment. The montage shows consecutive images from the area shown by the white square; recruitment of Dynamin is observed at 5:45 min (white arrow). B. The top graph represents the speed of the VLP shown in (A); the bottom graph demonstrates the dynamics of the intensity of both channels; the kymograph represents the change in intensity and position of the particle. C. Distribution of the number of dynamin recruitment events to bound VLP^Wu^:M^Ch^, n = 41. D. Distribution of time intervals between VLP^Wu^:M^Ch^ binding and the first recruitment of Dynamin, n = 37. E. Distribution of the time intervals between VLP^Wu^:M^Ch^ binding and VLP speed increase, n = 18. F. Cells treated with Dynole 34-2, which inhibits vesicle-mediated endocytosis, showing the inability of VLPs to enter cells. G. Cells expressing GFP-tagged Rab-5 showing lack of co-localization of the Rab5-positive vesicles with the VLP^Wu^:M^Ch^. H. Cells treated with LysoTracker showing co-localization of the VLP^Wu^:M^Ch^ with acidic vesicles (lysosomes or late endosomes).

Dynamin accumulation does not always reflect successful scission of the endocytic vesicles, and several abortive cycles are often observed before a productive scission occurs^42^. If a single VLP colocalizes transiently with Dynamin more than once, this would be indicative of such abortive events. Indeed, we observed that 64% of the particles colocalized with a Dynamin focus only once, 17% twice, and 19% experienced three or more colocalization events. These results suggest that in 36% of VLP entries there is at least one abortive Dynamin binding (Fig. 2C). In addition, Dynamin foci formation allowed us to measure the time between VLP binding to the membrane and endosome vesicle scission, which was 5.24±6.8 min. SPARTACUSS also allowed us to measure the time between Dynamin foci formation and VLP speed increase. Particles increased their speed within 45s of Dynamin binding, and, in 30% of cases, this increase occurred in parallel with the Dynamin scission event (Fig. 2D, E).

To further evaluate the role of endocytosis in VLP internalization, we used Dynol 34-2, a potent inhibitor of Dynamin 1, which prevents receptor-mediated endocytosis^41^. After Dynol 34-2 treatment, VLPs did bind to the cell surface but did not enter (Fig. 2F, Video S13). However, it should be noted that Dynol 34-2 also greatly altered cell morphology, inducing a round phenotype, indicative of an effect on the cell cortex. Such considerable changes in morphology could affect membrane-related processes, including endocytosis. To check if VLPs localized within the early endosome, we used cells expressing Rab5a-GFP. Surprisingly, we did not detect colocalization of the VLPs with early endosomes (Fig. 2G and video S14). As endosome maturation is paralleled by a drop in pH, we asked whether the VLPs colocalize with acidic vesicles^43^. To this end, we stained cells with Lysotracker, which marks acidic vesicles, after incubation of cells with VLPs. Some of the VLPs colocalized with the stained acidic vesicles (Fig. 2H and Video S15).

Taken together, our results indicate that Dynamin-mediated endocytosis is involved in VLP internalization. Furthermore, VLPs are not internalized via Rab5a-positive early endosomes but rather localize within acidic vesicles.

### Dynamics of VLP acidification

The pH of medium was previously shown to influence the internalization of an SARS-CoV-2 S protein-containing VSV chimera, with a more acidic environment promoting internalization preferentially via fusion ^44–46^. To evaluate the role of pH in VLP internalization, we sought to precisely measure the pH dynamics of VLPs in which a fraction of the M protein is tagged with superecliptic pHluorin in its C-terminal end ^47^. This protein emits bright fluorescent light in pH≥8 but its intensity sharply decreases at pH<7.5, completely disappearing at pH<5^48^. As the M protein C-terminal domain lies inside the VLPs, the fused pHluorin serves as a real-time indicator of the intra-VLP pH. The pHluorin-labelled VLPs in the medium (pH8) emitted a bright green signal; however, shortly after binding to the membrane of Vero E6 cells, the intensity rapidly disappeared (Video S16), suggesting that the pH of VLPs decreases sharply. To understand when this happens relative to VLP internalization, we used VLPs, which contain unlabelled E, S, N, and M proteins but also two labelled fractions of M protein - one with mCherry and another with pHluorin, in addition to the or cis-acting RNA element^8^. We will refer to these VLPs as VLP^Wu^:M^Ch^M^pH^ (short for VLP^Wuhan^:(E, S, N, M&M-mCherry&M-pHluorin). The dual VLP labelling enabled us to follow the VLP and analyze its dynamics even when complete disappearance of the pHluorin signal occurred due to a sharp drop in pH. We observed that in 63% of the VLPs, the pHluorin signal disappeared without any change in mCherry intensity, while in 34% of the VLPs both fluorescent signals disappeared simultaneously, and in 3% of the particles there was no change in the fluorescent signal during the time course of our experiment in Vero E6 cells (Fig. 3А). These results indicate that in 2/3 of the cases disappearance of the pHluorin fluorescent signal is a result of pH decrease without VLP disassembly.

**Fig. 3.**
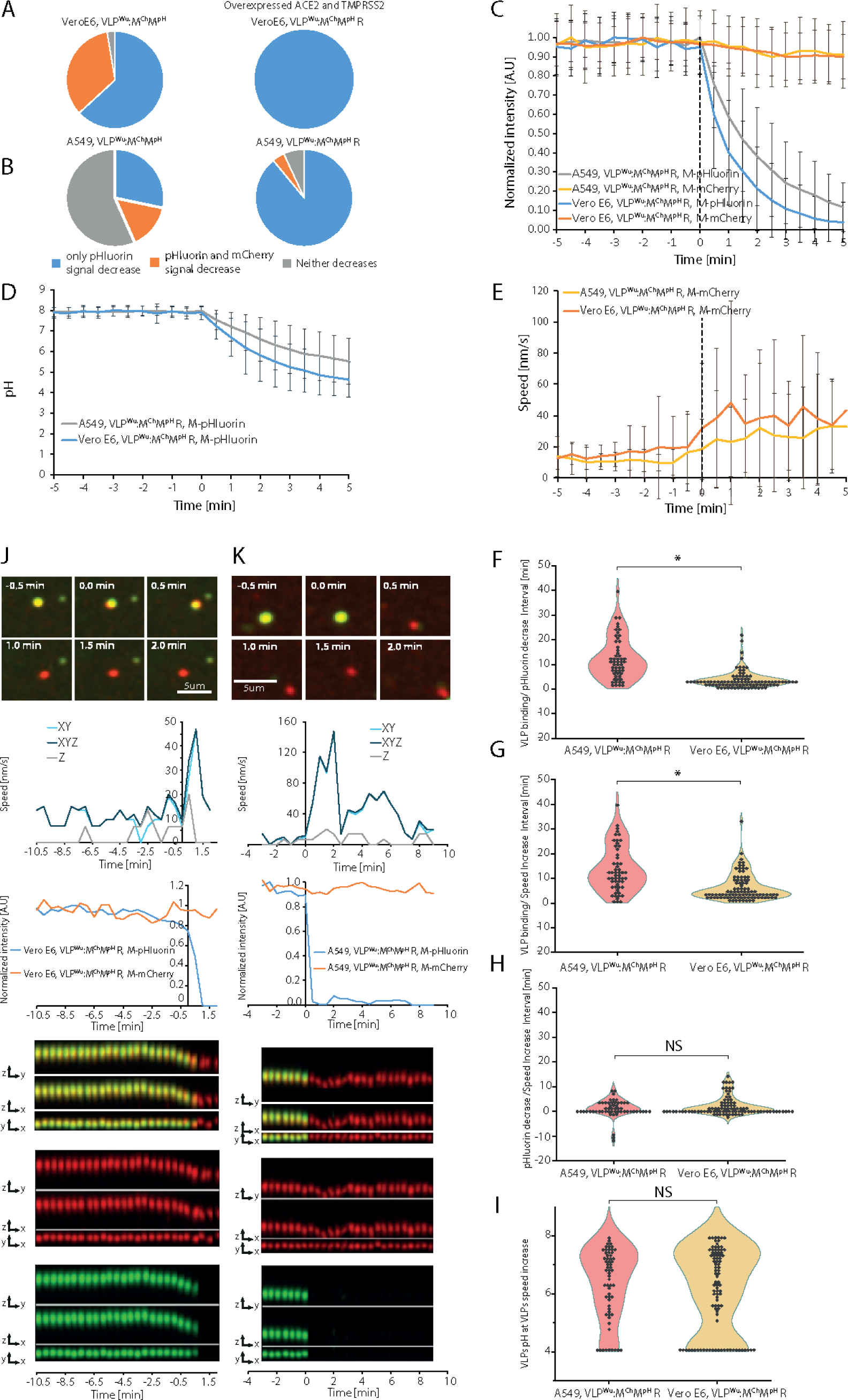
Dynamics of SARS-CoV-2 VLP^Wu^:M^Ch^M^pH^R binding, pH decrease, and speed increase in A549 and Vero E6 cells. A. Percentage of VLPs in which only M-pHluorin intensity decreases (Blue), the intensities of M-pHluorin and M-mCherry decrease simultaneously (Orange), or neither decreases (Gray) for 100 min after addition of VLP^Wu^:M^Ch^M^pH^R in VeroE6 cells with or without ACE2 and TMPRSS2 overexpression. B. Same experiment as (A) in A549 cells. C. Comparison of the M-pHluorin intensity decrease during internalization of VLP^Wu^:M^Ch^M^pH^R in A549 and Vero E6 cells. The average intensity of pHluorin is represented as a function of time where the individual VLPs were aligned to the start of VLP pHluorin decrease (0 min). The average M-mCherry intensity of the same particles is also presented. Errors bars represent the standard deviation. For A549 n = 55, for Vero E6 n = 93. D. Comparison of the dynamics of pH decrease during internalization of VLP^Wu^:M^Ch^M^pH^R in A549 and Vero E6 cells. The pH decrease is calculated based on the measured M-pHluorin intensity decrease. The average pH of VLPs is represented as a function of time where individual VLPs are aligned to the start of VLP pHluorin decrease (0 min). Error bars represent the standard deviation. For A549 n = 55, for Vero E6 n = 93. E. The average speed of VLP^Wu^:M^Ch^M^pH^R in A549 and Vero E6 cells measured based on the tracked M-mCherry signal. The average speed of VLPs was calculated after alignment of the individual VLP speeds to the start of the VLP pHluorin signal decrease (0 min). Error bars represent the standard deviation. For A549 n=55, for Vero E6 n=93. F. Distribution of time intervals between VLP^Wu^:M^Ch^M^pH^R binding and the start of pHluorin intensity decrease in A549 cells or Vero E6 cells. Two-tailed Student’s t-test; NS p>0.01; * p<0.01. For A549 n=55, for Vero E6 n=93. G. Distribution of time intervals between VLP^Wu^:M^Ch^M^pH^R binding and the start of VLP speed increase in A549 and Vero E6 cells. Two-tailed Student’s t-test; NS p>0.01; * p<0.01. For A549 n= 55, for Vero E6 n= 93. H. Distribution of time intervals between the start of VLP^Wu^:M^Ch^M^pH^R pHluorin intensity decrease and the start of VLP speed increase in A549 and Vero E6 cells. Two-tailed Student’s t-test; NS p>0.01; * p<0.01. For A549 n=55, for Vero E6 n=93. I. Distribution of the estimated pH of VLP^Wu^:M^Ch^M^pH^R, calculated based on the pHluorin signal at the moment when VLP speed started to increase in A549 and Vero E6 cells. Two-tailed Student’s t-test; NS p>0.01; * p<0.01. For A549 n=55, for Vero E6 n=93. J. Representative time-lapse images (top), corresponding VLP speed and intensity graphs (middle), and kymographs (merged, M-mCherry, and M-pHluorin) in all dimensions (bottom) for a single VLP^Wu^:M^Ch^M^pH^R undergoing internalization in an VeroE6 cell. In this example the speed increases in parallel with pHluorin signal decrease. K. Same as (J) in a A549 cell.

Next, we studied the influence of concurrent ACE2 and TMPRSS2 protease overexpression on VLP internalization. To this end, we used VLPs containing RNA (VLP^Wu^:M^Ch^M^pH^, R), which included the cis-acting T20 element reported to enhance packaging^8^. The inclusion of this RNA element did not affect the percentages of VLPs, in which the pHluorin signal disappeared without any change in mCherry intensity. Meanwhile, ACE2 and TMPRSS2 overexpression in cells led to a significant increase in the number of such VLPs (100%). Next, we performed the same experiments with human lung adenocarcinoma A549 cells, which are the standard pulmonary epithelial cell model for SARS-CoV-2 infection (Video S17). Without ACE2 and TMPRSS2 overexpression, 28% of the VLPs exhibited a decrease in pHluorin fluorescence without any change in mCherry intensity, 15% exhibited simultaneous disappearance of both signals, and 57% did not show any change in both signals throughout the experiment (Fig. 3B). These results suggest that A549 cells are less susceptible to SARS-CoV-2 VLP internalization than Vero E6 cells. Overexpression of ACE2 and TMPRSS2 considerably increased the fraction of VLPs exhibiting a decrease in pHluorin intensity, without changes in mCherry intensity (89% versus 57%). Overall, ACE2 and TMPRSS2 overexpression increased the number of VLPs exhibiting a decrease in pH without VLP disassembly in both cell lines (Fig 3A, B).

To evaluate the speed with which the pHluorin signal decreases, we aligned all VLP tracks to the start of the pHluorin intensity decrease (Fig. 3C). We thus measured the half-time of pHluorin signal disappearance which was 1.4 and 1.6 min in Vero E6 and A549 cells, respectively. Conversion of pHLuorin intensity to pH values ^47^ showed that the VLP pH in Vero E6 cells decreased from 8 to 6.3, while that in A549 cells decreased from 8 to 6.9 over a period of 1.5 min (Fig. 3C, D), attesting to the rapid acidification of VLPs. Our results also demonstrated that the speed of VLPs tends to increase following the start of pH decrease (Fig. 3-E).

Dual labelling with both M-mCherry and M-pHluorin allowed us to infer the temporal order of the three stages of VLP internalization at the single VLP level: VLP binding to the cell membrane, VLP pH decrease, and the increase of its speed. We thus measured the time intervals between each possible pair of the above-described internalization steps (Fig. 3F, G, H, and Table 1). In VeroE6 cells, VLP pH began to decrease 4.1 ± 3.6 min after plasma membrane binding, and 2.4 ± 3.7 min later the VLP speed started to increase (Table 1). In A549 cells, VLP pH began to decrease 12.5 ± 8.4 min after binding, and 1.4 ± 3.7 min later the VLP speed started to increase. These results indicate that, on average, the pH change occurs during or immediately before the VLP speed increases (Fig. 3J, K). At a single VLP level, however, we have examples where the speed increase occurs after the pH decreases, but also examples where the speed increases simultaneously or before the start of pH decrease (Extended Data Fig. 3 and 4, and Video S18-S21)), indicating that there is no direct causal relationship between the two events. This uncoupling is further supported by the significant variation in pH at which the VLP speed increase begins (Fig. 3I). As we demonstrate above, the speed increase is a hallmark for active microtubule movement of the vesicles containing VLPs. Taken together, for the majority of VLPs, pH acidification starts before or during microtubule attachment (Extended Data Fig 3. C) As acidification takes less than 2 min, we observe cases when movement via the microtubules occurs after the pH is already <5. Direct comparison of VLP internalization kinetics among the two cell lines (Fig. 3H) revealed no statistically significant difference in the interval between VLP pH decrease and speed increase (start of the microtubule movement). However, the intervals between VLP binding and both pH decrease and speed increase are 2-3 times shorter in VeroE6 cells than in A549 cells (Fig. 3, Table 1). This difference in internalization efficiency may contribute to the greater SARS-CoV-2 susceptibility of Vero E6 relative to A549 cells, generally attributed to the lack of interferon signalling in the former^49–52^. Close examination of the distribution of the intervals between VLP binding and both pH decrease and speed increase in A549 cells revealed a small population of VLPs for which these intervals are significantly longer (Fig. 3F, G). These longer intervals may be attributed to cycles of abortive dynamin-mediated endosome scission, as previously discussed.

**Table 1.** Measured intervals between different steps of VLP internalization.

| VLP Variants/cell line | VLP binding to pH drop or N release | pH drop or N release to speed increase | VLP binding to speed increase |
| --- | --- | --- | --- |
| VLP-WT MM in Vero E6 | 4.1 ± 3.6 min | 2.4 ± 3.7 min | 6.5 ± 5.4 min |
| VLP-WT MM in A549 | 12.5 ± 8.4 min | 1.4 ± 3.7 min | 13.9 ± 9.2 min |
| VLP-WT NM in A549 | 13 ± 8.6 min | 7.0 ± 10.0 min | 20.1 ± 14.8 min |
| VLP-Omi NM in A549 | 14.2 ± 8.5 min | 2.9 ± 6.9 min | 17.1±10.1 min |
| VLP-del1 MM in VeroE6 | 6.4 ± 5.4 min | 2.8 ± 8.2 min | 9.2 ± 8.9 min |
| VLP-del1 MM in A549 | 16.7 ± 12.7 min | 1.7 ± 1.6 min | 18.4 ± 12.9 min |
| VLP-del1 MM in A549_381_New | 10.12 ± 8.72 min | 3.61 ± 7.79 min | 13.74 ± 11.57 min |
| VLP-OMI MM in A549 | 13.01 ± 8.4 min | 4.03 ± 7.44 min | 17.04 ± 11.48 min |

In summary, our findings highlight the notable speed and efficacy of VLP internalization, which are more pronounced in VeroE6 relative to A549 cells. Furthermore, the dynamic profiles of VLP internalization-related processes, namely, membrane binding, acidification, and initiation of microtubule transport, show that the latter two processes, while temporally proximal, are not interdependent.

### Kinetics of VLPs lacking the furin cleavage site

In contrast to SARS-CoV, SARS-CoV-2 harbors a PRRA furin cleavage site (FCS) at the S1/S2 junction of the S protein^52^. Cleavage by cellular protease furin followed cleavage at the S2’ site by TMPRSS2 or cathepsins results in the separation of the two S sub-domains. Thus, we sought to determine how absence of the FCS would affect internalization at the single VLP level. To this end, we used VLPs containing M-mCherry, M-pHluorin, and S protein lacking an FCS (del-1)^53–57^. We refer to these as VLP^Wu(del-1)^:M^Ch^M^pH^,R, short for VLP^Wuhan^ ^(del-1)^:(N, E, S, M&M-mCherry & M-pHluorin, T20 RNA). Treatment of A549 cells overexpressing ACE2 and TMPRSS2 with VLP^Wu(del-1)^: M^Ch^M^pH^ revealed a decrease in the percentage of VLPs for which pHluorin signal decrease occurred without a change in mCherry intensity, from 89% for VLP^Wu^:M^Ch^M^pH^, R to 73% for VLP^Wu(del-1)^:M^Ch^M^pH^, R (Fig. 4A). Further, the rate of pHluorin decrease was similar between the two (Fig. 4B). The distribution of the time intervals between VLP binding and pHluorin decrease/VLP speed increase was also comparable (Fig. 4C-G, Table 1). As observed for VLP^Wu^, the pH decrease of VLP^Wu(del-1)^ initiated either a little before, in parallel to, or after the speed of the same VLPs began to increase (Extended Data Fig. 5, and Video S22, S23). Similar results were obtained in VeroE6 cells (Extended Data Fig. 6 and 7 Video S24, S25).

**Fig. 4.**
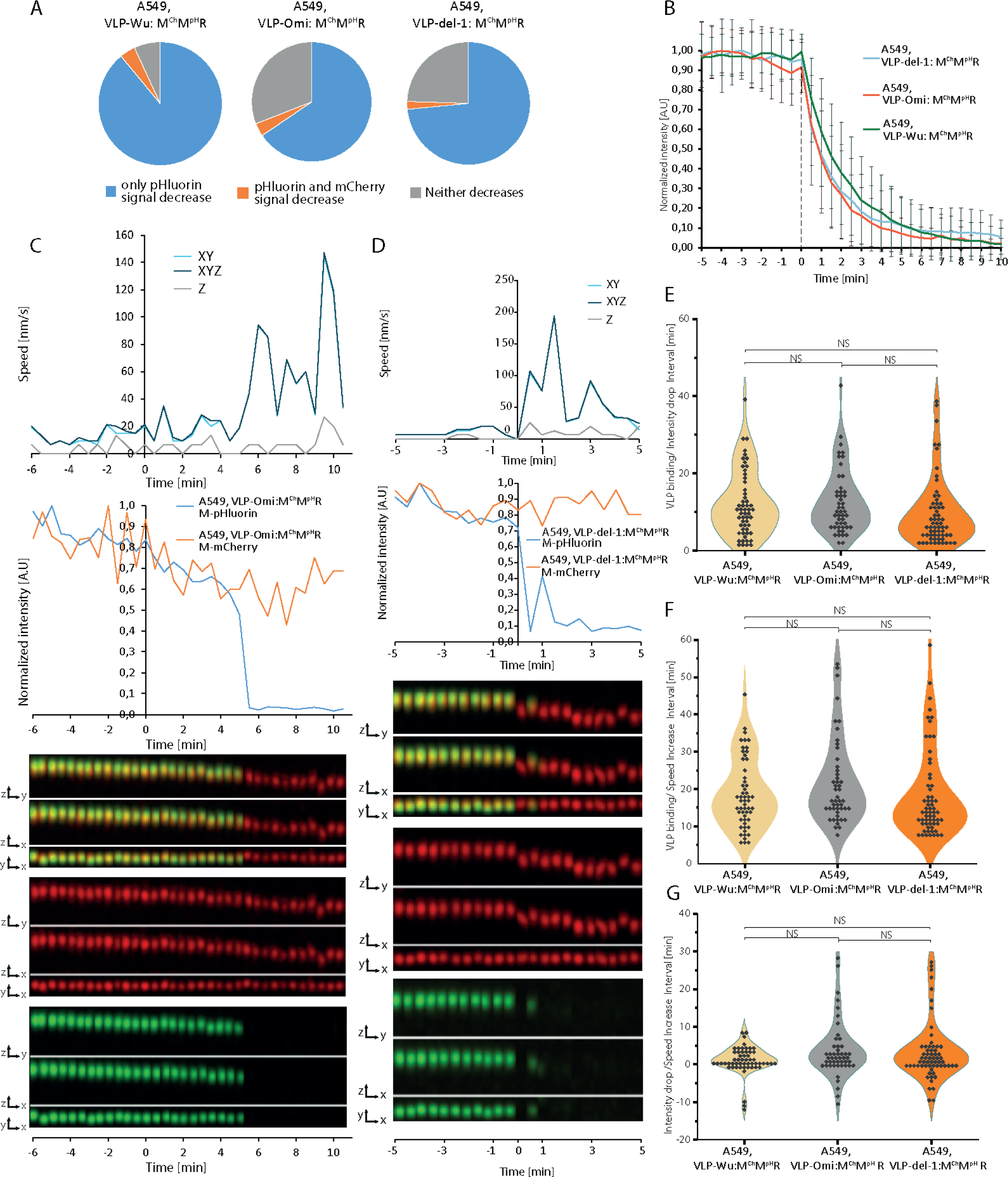
Comparison of VLP binding, acidification, and speed increase dynamics between VLP^Wu^: M^Ch^M^pH^R, VLP^Omi^: M^Ch^M^pH^R and VLP^del-1^: M^Ch^M^pH^R during internalization in A549 cells. A. Percentage of VLPs in which only the M-pHluorin intensity decreases (Blue), M-pHluorin and M-mCherry intensities decrease simultaneously (Orange), or neither decreases (Gray) B. Comparison of pHluorin intensity decrease during the internalization of VLP^Wu^: M^Ch^M^pH^R, VLP^Omi^: M^Ch^M^pH^R and VLP^del-1^: M^Ch^M^pH^R in A549 cells. The average intensity of pHluorin is represented as a function of time where the individual VLPs were aligned to the start of VLP pHluorin decrease (0 min). The average M-mCherry intensity is also presented. Error bars represent the standard deviation. For VLP^Wu^: M^Ch^M^pH^R n=55, for VLP^Omi^: M^Ch^M^pH^R n=48, for VLP^del-1^: M^Ch^M^pH^R n=62. C. Representative VLP speed and intensity graphs (top) and kymographs (merged, M-mCherry, and M-pHluorin) in all dimensions (bottom) for a single VLP^Omi^: M^Ch^M^pH^R undergoing internalization in an A549 cell. In the example the speed increases in parallel with pHluorin signal decrease. D. Same as (C) but for VLP^del-1^: M^Ch^M^pH^R. E. Distribution of time intervals between VLP binding and start of pHluorin intensity decrease for individual VLP^Wu^: M^Ch^M^pH^R, VLP^Omi^: M^Ch^M^pH^R, and VLP^del-1^: M^Ch^M^pH^R during internalization in A549 cells. Two-tailed Student’s t-test; NS p>0.01; * p<0.01. VLP^Wu^: M^Ch^M^pH^R n=55, for VLP^Omi^: M^Ch^M^pH^R n=48, for VLP^del-1^: M^Ch^M^pH^R n=62. F. Distribution of time intervals between VLP binding and start of speed increase for individual VLP^Wu^: M^Ch^M^pH^R, VLP^Omi^: M^Ch^M^pH^R, and VLP^del-1^: M^Ch^M^pH^R during internalization in A549 cells. Two-tailed Student’s t-test; NS p>0.01; * p<0.01. For А549-WT n=55, for A549-Omi n=48, for А549-del1 n=62. G. Distribution of time intervals between VLP intensity decrease and start of speed increase for individual VLP^Wu^: M^Ch^M^pH^R, VLP^Omi^: M^Ch^M^pH^R, and VLP^del-1^: M^Ch^M^pH^R during internalization in A549 cells. Two-tailed Student’s t-test; NS p>0.01; * p<0.01. For VLP^Wu^: M^Ch^M^pH^R n=55, for VLP^Omi^: M^Ch^M^pH^R n=48, VLP^del-1^: M^Ch^M^pH^R n=62.

Taken together, our results indicate no significant role for the FCS in the internalization of SARS-CoV-2 VLPs. It is known that passage of SARS-CoV-2 in VeroE6 cells leads to mutations in the S1/S2 junction, suggesting that the FCS is dispensable for virus propagation, in line with our results^57,58^. The lack of significant difference in internalization dynamics between VLPs with and without FCS deletion observed herein may attributed to non-effective cleavage by furin during maturation even in VLPs with an intact FC.

### Kinetics of Omicron VLPs

Emergence of the SARS-CoV-2 Omicron variant in the autumn of 2021 was followed by its rapid spread, overtaking previous variants in global prevalence. Omicron harbors more than 50 amino acid substitutions, 37 of which in the S protein, with 15 affecting the receptor-binding domain. In light of its considerable transmissibility and rapid replication in human bronchi (70-fold greater than of previous variants), we sought to measure the internalization kinetics of Omicron^59^. To this end, we employed VLPs composed of unlabelled N, E, S, and M proteins harboring Omicron substitutions. VLPs also included mCherry-tagged and pHluorin-tagged M protein and are referred to as VLP^Omi^:M^Ch^M^pH^R, short for VLP^Omicron^:(N, E, S, M, M-mCherry & M-pHluorin, T20 RNA). Treatment of A549 cells overexpressing ACE2 and TMPRSS2 (Video S26, S27, S28) revealed a decrease in the proportion of VLPs in which the pHluorin signal decay occurred without a change in mCherry signal, that is, from 89% in VLP^Wu^:M^Ch^M^pH^R to 65% for VLP^Omi^:M^Ch^M^pH^R (Fig. 4A). However, the rate of decrease pHluorin intensity was largely identical between the two VLP types as well as the Wuhan del-1 mutant. The average and the distribution of intervals between VLP binding and pHluorin decrease/VLP speed increase were also comparable among the three VLP types (Fig. 4E, Extended Data Fig. 8). Taken together, these data suggest that, despite the considerable number of S mutations and the greater transmissibility observed in humans, no considerable differences in internalization kinetics were noted for the Omicron variant. Thus, our findings point away from enhanced internalization as a basis for superior transmissibility, suggesting greater stability or replication efficiency as potential underlying mechanisms.

### Rate of VLP nucleocapsid release

Nucleocapsid release occurs via VLP fusion either to the plasma or endosomal membrane, representing an essential step in the internalization process. Thus, we set out to measure the dynamics of VLP nucleocapsid release, using VLPs in which a fraction of the nucleocapsid-forming N protein is tagged with EGFP, and a fraction of M is tagged with mCherry (VLP^Wu^:N^eG^M^Ch^R), short for SARS-CoV-2:(E, S, N&N-eGFP, M&M-mCherry, T20 RNA) VLPs. We observed that, while initially greater than 90% of VLPs emitted both green and red fluorescence less than 1% of VLPs emitted green fluorescence at 2 days after production, which would suggest that tagging N strongly reduces VLP stability, as previously reported^41^. In 57% of these double-labelled VLPs, the EGFP signal disappeared before the mCherry one, which may reflect N protein release (Fig. 5A). The rate of N-EGFP signal decay in VLP^Wu^:N^eG^M^Ch^R was slower than that of M-pHluorin intensity decrease in VLP^Wu^:M^Ch^M^pH^R (Fig. 5C). The observed slow decay suggests that nucleocapsid release may not be a single-step rapid event, but a rather gradual process of continuous N-EGFP release, which cannot be followed thereafter. Only in a single case (of 18 VLPs) were we able to follow the N-eGFP signal after rapid release of the nucleocapsid. We tracked both fluorescent signals (M-mCherry and N-eGFP) for this VLP and observed that the M-mCherry signal did not move in z during signal separation, while the eGFP did. This could suggest that in this case fusion occurs either at the cell plasma membrane or during VLP ingression, prior to active movement via the microtubular network (Fig. 6 and Video S29). On average, there was no statistically significant difference between the interval from VLP binding to the start of pH decrease of VLP^Wu^:M^Ch^M^pH^R (Table 1) and the interval from VLP binding to the start of nucleocapsid release of VLP^Wu^:N^eG^M^Ch^R (Fig. 5D-F). This result suggests that, on average, VLP acidification coincides with nucleocapsid release.

**Fig. 5.**
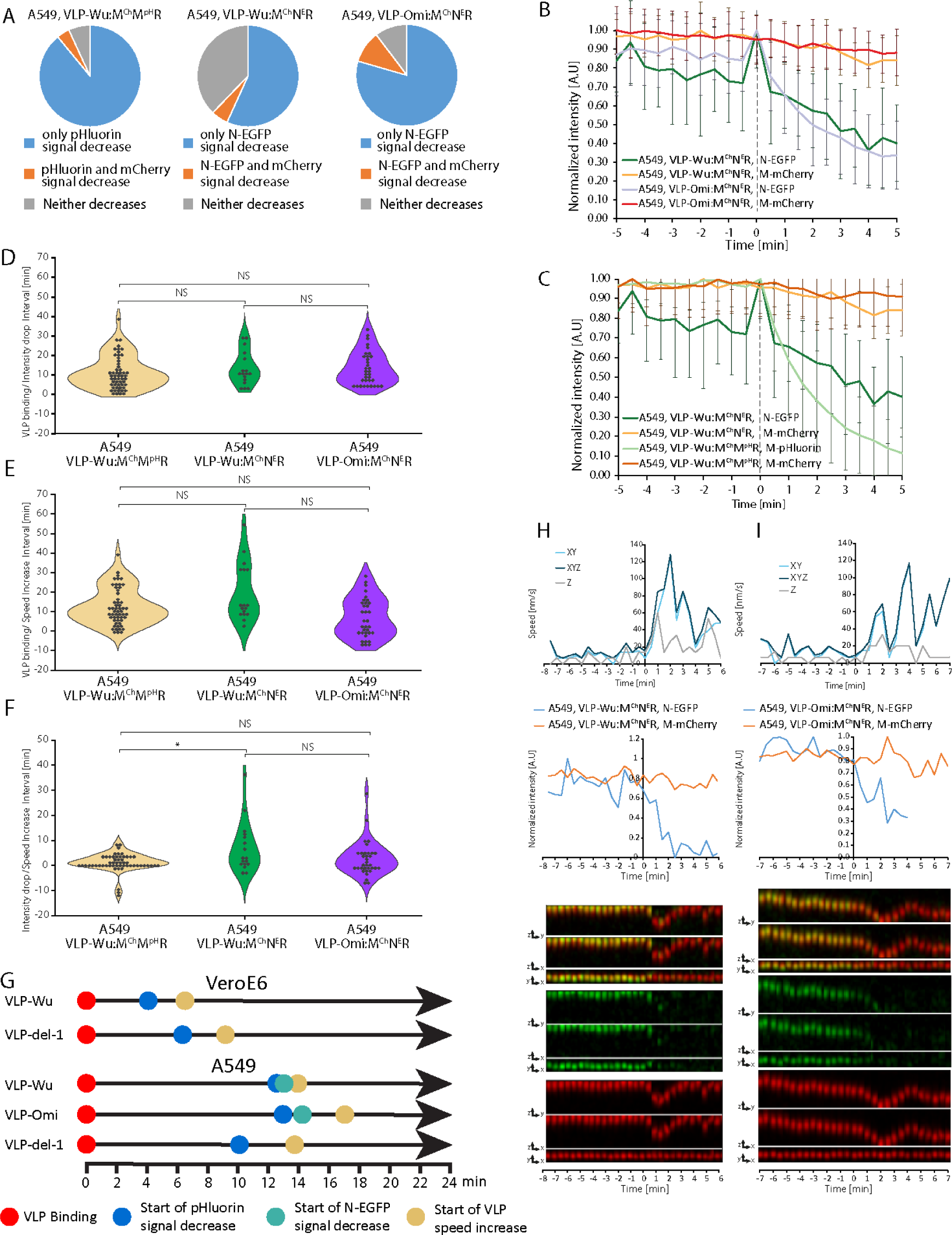
Comparison of VLP binding, acidification, and speed increase dynamics between VLP^Omi^:M^Ch^N^E^R, VLP^Wu^:M^Ch^N^E^R, and VLP^Wu^:M^Ch^M^pH^R during internalization in A549 cells. A. Percentage of VLPs in which only the signal intensity of N-EGFP (middle and right) or M-pHluorin (left) decreases (Blue), the intensities of both N-EGFP/M-pHluorin and M-mCherry decrease simultaneously (Orange), and neither decreases (Gray). VLPs were tracked for 100 min after addition. B. Comparison of N-EGFP intensity decrease rate during VLP^Omi^:M^Ch^N^E^ R and VLP^Wu^:M^Ch^N^E^R internalization in A549 cells. The average intensity of N-EGFP is presented as a function of time where the individual VLPs were aligned to the start of N-EGFP signal decrease (0 min). The average M-mCherry intensity is also presented. Error bars represent the standard deviation. For A549 VLP^Wu^:M^Ch^N^E^R n=17, for A549 VLP^Omi^:M^Ch^N^E^ n=34. C. Comparison of M-pHluorin and N-EGFP intensity decrease rates during internalization of VLP^Wu^:M^Ch^M^pH^R and VLP^Wu^:M^Ch^N^E^R, respectively, in A549 cells. The average intensities of N-EGFP and M-pHluorin are presented as a function of time where the individual VLPs were aligned to the start of VLP N-EGFP or M-pHluorin decrease (0 min). The average M-mCherry intensity of the same particles is also presented. Error bars represent the standard deviation. For A549 VLP^Wu^:M^Ch^M^pH^R n=55, for A549 VLP^Wu^:M^Ch^N^E^R n=17. D. Distribution of time intervals between VLP binding and the start of pHluorin/N-EGFP intensity decrease for individual VLP^Wu^:M^Ch^M^pH^R, VLP^Wu^:M^Ch^N^E^R, and VLP^Omi^:M^Ch^N^E^R during internalization in A549 cells. Two-tailed Student’s t-test; NS p>0.01; * p<0.01. For A549 VLP^Wu^:M^Ch^N^E^R n=17, for VLP^Omi^:M^Ch^N^E^R n=34 and VLP^Wu^:M^Ch^M^pH^R n=55. E. Distribution of time intervals between VLP binding and start of speed increase for individual VLP^Wu^:M^Ch^M^pH^R, VLP^Wu^:M^Ch^N^E^R, and VLP^Omi^:M^Ch^N^E^R during internalization in A549 cells. Two-tailed Student’s t-test; NS p>0.01; * p<0.01. For A549 VLP^Wu^:M^Ch^N^E^R n=17, for A549 VLP^Omi^:M^Ch^N^E^R n=34 and A549 VLP^Wu^:M^Ch^M^pH^R n=55. F. Distribution of time intervals between start of pHluorin/N-EGFP intensity decrease and start of speed increase for individual VLP^Wu^:M^Ch^M^pH^R, VLP^Wu^:M^Ch^N^E^R, and VLP^Omi^:M^Ch^N^E^R during internalization in A549 cells. Two-tailed Student’s t-test; NS p>0.01; * p<0.01. For A549 VLP^Wu^:M^Ch^N^E^R n=17, for A549 VLP^Omi^:M^Ch^N^E^R n=34 and A549 VLP^Wu^:M^Ch^M^pH^R n=55. G. Schematic of the major internalization-associated events through time for wild-type and mutant VLPs in VeroE6 and A549 cells. H. Representative VLP speed and intensity graphs (top) and kymographs (merged, M-mCherry, and N-EGFP) in all dimensions (bottom) for a single VLP^Wu^:M^Ch^N^E^R undergoing internalization in an A549 cell. In this example the speed increases in parallel with pHluorin signal decrease. I. Same as (H) for VLP^Omi^:M^Ch^N^E^R.

**Fig. 6.**
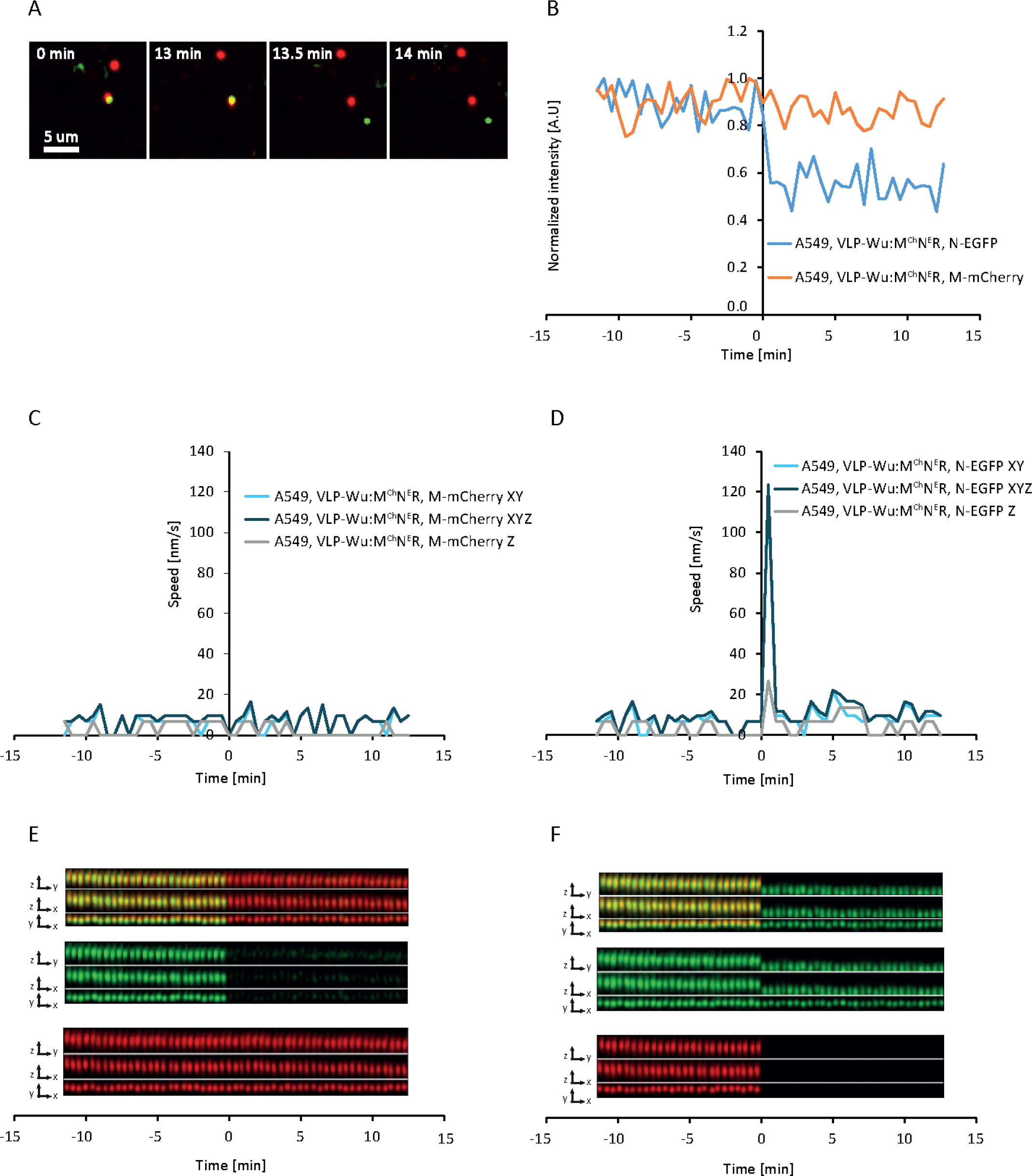
Tracking of nucleocapsid release following VLP internalization. A. Time-lapse images of N-EGFP release during internalization of VLP^Wu^:M^Ch^N^E^R in A549 cells where the separation of the nucleocapsid (N-EGFP) from the VLP membrane (M-mCherry) can be observed. B. Changes in N-EGFP and M-mCherry during nucleocapsid release for the above VLP (tracked based on the N-EGFP signal). C. VLP speed profile during nucleocapsid release measured based on M-mCherry movement for the same VLP. D. VLP speed profile during nucleocapsid release measured based on N-EGFP movement for the same VLP. Note the speed increase during nucleocapsid release, which is missing in (C). E. Kymogram in all dimensions measured based on M-mCherry tracking. F. Kymogram in all dimensions based on N-EGFP tracking. Note the change in movement along the Z-axis during nucleocapsid release, which is missing in (E).

Next, we sought to determine nucleocapsid release kinetics in omicron VLP^Omi^:M^Ch^N^eE^R, short for SARS-CoV-2^Omicron^:(E, S, M&M-mCherry, N&N-eGFP, T20 RNA) VLPs. Surprisingly, the percentage of intact, double-labelled particles was considerably higher among VLP^Omi^:M^Ch^N^eE^R (2-3%) as opposed to their VLP^Wu^:M^Ch^N^eE^R counterparts (<1%), while remaining much lower than that among VLP^Omi^:M^Ch^M^pH^R (>90%). In approximately 80% of VLP^Omi^:M^Ch^N^eE^R, N-eGFP fluorescence disappeared before mCherry, reflecting a 23% more effective release than observed for VLP^Wu^:M^Ch^N^eE^R (Fig. 5A). However, the rates of N-eGFP signal decrease (Fig. 5B) were similar between the Wuhan and Omicron VLPs, which suggests that nucleocapsid release occurred at a comparable rate. In addition, there was no statistically significant difference between the interval from VLP binding to nucleocapsid release for the Wuhan and Omicron VLPs (Fig. 5D, Table 1). Thus, omicron mutations had no effect on the nucleocapsid release step of internalization. In the Omicron VLP, speed increase occurred shortly after nucleocapsid release, as observed for Wuhan VLP (Fig. 5E, F, Table 1) indicating that VLP membrane fusion occurs, on average, before the start of microtubule-mediated transport. However, at single VLP level, we observed cases of speed increase occurring after, in parallel to, or before nucleocapsid release (Fig. 5H, I and Extended Data Fig. 9 and 10, and Video S30-33), indicating no causality between these two events, as was also observed above for the pH decrease.

In this work we developed a customised software pipeline (SPARTACUSS) to reconstruct the timeline defined by five of the hallmark parameters for viral internalisation on a single SARS-CoV-2 VLP level - VLP binding, start of pH decrease, start of nucleocapsid release, Dynamin-VLP co-localization and start of active microtubule dependent VLP movement (Fig. 5G, Table 1). Comparison between the timing of these events demonstrates that about 4 minutes after VLP binding to Vero E6 cells and 12 minutes after VLP binding to A549 cells, the pH starts to rapidly decrease which coincides with Dynamin binding and nucleocapsid release quickly followed by initiation of active microtubule dependent VLP motion. Our results suggest that VLP fusion (N-nucleocapsid release) occurs simultaneously with or shortly after endosome formation. Surprisingly, the VLPs do not colocalize with Rab-5a positive early endosomes during VLP internalization. The two to three times shorter time required for VLP internalization in VeroE6 cells compared to A549 cells could explain VeroE6 higher susceptibility to SARS-CoV-2 infection. Neither Omicron nor del-1 mutations influence the internalization steps of SARS-CoV-2 VLPs. Representative time-lapse videos of all investigated internalization events in different cell lines can be accessed in the dedicated [COVIDynamics database](https://covidynamics.imb.bas.bg/). It is our vision that the comprehensive measurement of the SARS-CoV-2 VLP internalization steps will facilitate the study of the molecular mechanisms of the SARS-CoV-2 life cycle on a single-particle level as well as the evaluation of antiviral therapeutics.

## Materials and Methods

### Cell lines and transfections

The VeroE6, U2OS, and A549 cells used in this study were obtained from American Type Cell Culture (ATCC). VeroE6-ACE2-TMPRSS2 (VeroE6-AT) and A549-ACE2 clone 8-TMPRSS2 (A549-AT) cells, which overexpress ACE2 and TMPRSS2, were provided by the NIBSC Research Reagent Repository, UK (catalogue numbers 101003 and 101006, respectively) and were a kind gift from Prof. Arvind Patel, University of Glasgow. VeroE6 and U2OS cell lines were cultured in Dulbecco’s Modified Eagle Medium (DMEM, Thermo Fisher Scientific, Waltham, MA, USA) with a high content of glucose, 10% fetal bovine serum (FBS), and 100 units/mL penicillin and 100 μg/mL streptomycin at 37 °C and 5% CO_2_. A549 cell lines were cultured in Roswell Park Memorial Institute medium (RPMI, Gibco) supplemented with 10% (FBS) 100 units/mL penicillin and 100 μg/mL streptomycin at 37 °C and 5% CO2. In addition, 2 mg/mL Geneticin and 200 µg/mL Hygromycin B were added to the culture medium of VeroE6-AT and A549-AT cells, respectively.

### Transfection

For expression of Ace2NeonGreen and mNeonGreen, we performed transient transfection via baculovirus-mediated gene transduction of mammalian cells (BacMam) using the Montana Molecular ACE2 green kit (product #C110G) as per the manufacturer’s protocol. Expression of the fluorescent proteins was evaluated via fluorescence microscopy two days after transfection, whereafter cells were treated with VLPs.

For expression of eGFP-tagged Dynamin 1, we used a plasmid that was a gift from Sandra Schmid (Addgene plasmid # 34680; http://n2t.net/addgene:34680; RRID: Addgene_34680)^27^. To inhibit the dynamin-mediated entry into cells we used Dynole® 34-2, a dynamin I/II inhibitor (ab120463, abcam). Cells were treated with 30 µmol Dynole for 15 min and were then imaged every 30 sec in 31 Z-planes, with a Z-step size of 0.2 µm.

To observe endosome trafficking, we used CellLight™ Early Endosomes-GFP, BacMam 2.0 (Catalog number: C10586, Invitrogen), which allowed us to introduce GFP-tagged Rab5a into cells and thus visualize early endosome vesicles. We plated 10 000 cells in 35-mm glass-bottom culture dishes (MatTek Corporation, Ashland, MA, USA) and incubated these with 2 µL of CellLight™ Early Endosomes-GFP overnight. On the following day, we added VLPs and imaged the cells at 30-s intervals in 11 Z-planes with a Z-step size of 0.2 µm.

To observe late endosomes, we used LysoTracker (ThermoFisher Scientific). Cells were incubated with the VLPs for 30 min, treated with 50 mM Lysotracker for 1 min, and imaged every 30 s in 11 Z-planes with a Z-step size of 0.2 µm.

To image tubulin, we used abberior LIVE 610 conjugated to cabazitaxel. Cells were inoculated with VLPs for 30 min and incubated with abberior LIVE 610 conjugated to cabazitaxel for 15 min, followed by washing with fresh FluoroBrite™ DMEM (Gibco). The cells were imaged every 30 s in a single Z-plane.

### Time-lapse live-cell imaging

Forty-eight h before imaging, all cells were transferred to MatTek glass-bottom dishes (MatTek Corporation, Ashland, MA, USA) at 20% confluence. Live-cell imaging was performed on an Andor Dragonfly spinning-disk confocal system with a Nikon Eclipse Ti2-E inverted microscope equipped with the Nikon Perfect Focus System (PFS), a Nikon CFI Plan Apo VC 60x (NA 1.2) water immersion objective, a Nikon Apo 60x (NA 1.4) oil objective or Nikon HP Plan Apo 100x (NA 1.35) silicone λS objective, and a high-sensitivity iXon 888 Ultra Electron Multiplying Charge-Coupled Device (EMCCD) camera. Time intervals between consecutive frames varied between 15 and 30 s depending on the type of experiment, cell line, and the labelled protein. Images were acquired with variable z stacks of between 1 and 36 steps depending on the type of experiment and a z-step size of 0.2 µm. Prior to imaging, Petri dishes mounted on the microscope were left to thermally equilibrate for at least 30 min. All cells were incubated in FluoroBrite™ DMEM (Gibco) for imaging and maintained at 37°C and 5% CO_2_ during imaging. To visualize cells as transparent and opaque, we used the Imaris imaging software tool (Oxford Instruments).

### Electron microscopy

MLE-12 cells (a gift from Dr Kristi Warren) were grown on ACLAR disks and incubated with SARS-CoV-2 VLP^Wu^ for 5 min. Cells were fixed in 2.5% glutaraldehyde plus 1% paraformaldehyde in 0.1 M cacodylic buffer for 30 min and then embedded in resin using an Embed 812 kit (Electron Microscopy Sciences, Hatfield, PA, USA) and sectioned at 80 nm with a diamond knife (Diatome) using a Leica EM UC6 (Leica Microsystems, Wetzlar, Germany). Sections were visualized using a JEM 1400 Plus electron microscope (JEOL, Tokyo, Japan) at 120 kV.

### Virus-like particle preparation

VLPs were prepared as previously described^11,12^. Once generated, these were kept on ice until use 1 to 6 days after production.

### Antibody inhibition of SASR-CoV-2 VLPs

We diluted VLPs in 25 µL of FluoroBrite™ DMEM (Gibco) without FBS and then added 2 µL of Recombinant Anti-SARS-CoV-2 Spike Glycoprotein S1 antibody [CR3022] (Abcam number-ab273073). After incubating this mixture for 30 min, we added it to cells and proceeded with image acquisition.

### VLP tracking

The channel containing VLP fluorescence was isolated in Fiji, smoothed with 3D Gaussian filtering (1.5-pixel radius in XY and Z. Maximum intensity projection (MIP) was then performed.

The particles were tracked using the MTrackJ plugin, clicking on the particle location in each time frame, using the option “Apply local cursor snapping during tracking” with a range of 5x5 pixels. Completed tracks were exported as .mdf files. Multiple particles were tracked in the same session. To continue previous tracking sessions, we would import the previous .mdf file and continue, so that one .mdf file contains all tracks of one movie.

The subsequent processing steps outlined below were performed with a set of python scripts and Fiji macros.

To convert the 2D+t tracks to 3D+t, the script adds the Z coordinate by going back to the 3D movie before MIP and finding the plane that contains the maximum signal along a cylinder with a 2-pixel radius, centered at the location of the particle.

Instantaneous VLP speed v(t) at each time t was measured by subtracting the positions of a particle at times (t+1) and t, then dividing by the time interval between successive frames. The positions were converted to real physical units (nanometers) using the pixel sizes in XY and Z.

To visualize single particles over time, a cuboid with size dx, dy, and dz was cropped out of the 3D image stack for each time point, such that the particle is located at the center of the square (dx, dy), while dz is equal to the full width of the stack. Thus, the motion of the particle in Z can be visualized. The cuboids were maximum-projected along either the X or Y axis, resulting in ZY or ZX rectangles, respectively. For completeness, we also generated the XY squares, by performing max-Z projections. The stacking of these three ZY, ZX, and XY crops is referred to as a kymograph and discussed throughout the paper.

### VLP analysis

After having information on the position, speed, and intensity of each particle, we determined the precise moment when the intensity started to decrease and the speed started to increase - specific hallmarks of the VLP internalization process. Every particle was then aligned to these positions, and an average for the speed and intensity was obtained. For every intensity value a measured background value was subtracted. The curves were then plotted and compared showing the standard deviation for each point as error bars.

When determining the pH of each particle we used the intensity values and transformed them to pH values using the following formula: 7.11-LOG10(1/(X*0.886)-1)^47^, where X stands for the corresponding intensity value.

### Statistical analysis

Between-group comparisons were performed using the Student’s t-test. The significance threshold was set at p<0.01. Data are presented as the mean ± standard deviation.

## Acknowledgements

This research was funded by: NSF 2026657 (MV and SS), NSF 2102948 to (MV and SS) and NIH R56 AI150474-06A1 to (SS and MV), Ministry of Education and Science ДО1-166 (SSS, AA, AI and RS). We thank Petar-Bogomil Kanev for critically reading the manuscript.

## Author contributions

SSS, SS. and MV designed the experiments. SSS, AA, AI, performed microscopy experiments. W.P., R.G., H.D performed virus-like particle preparation and characterization, AA and AI conducted the data analysis. RS developed the software tools. SSS, SS, RS and AA wrote the manuscript. All authors reviewed and edited the text.

## Data Availability

To make the results publicly available we generated the COVIDynamics database with representative videos for all the experiments at COVIDynamics.imb.bas.bg. The database is searchable and accessible. As the raw data is more than 100 TB it would be provided upon reasonable request. The SPARTACUSS software will be open-access with detailed, step-wise manual and everybody will be able to use it.

## Extended Data

**Extended Data Fig.1.**
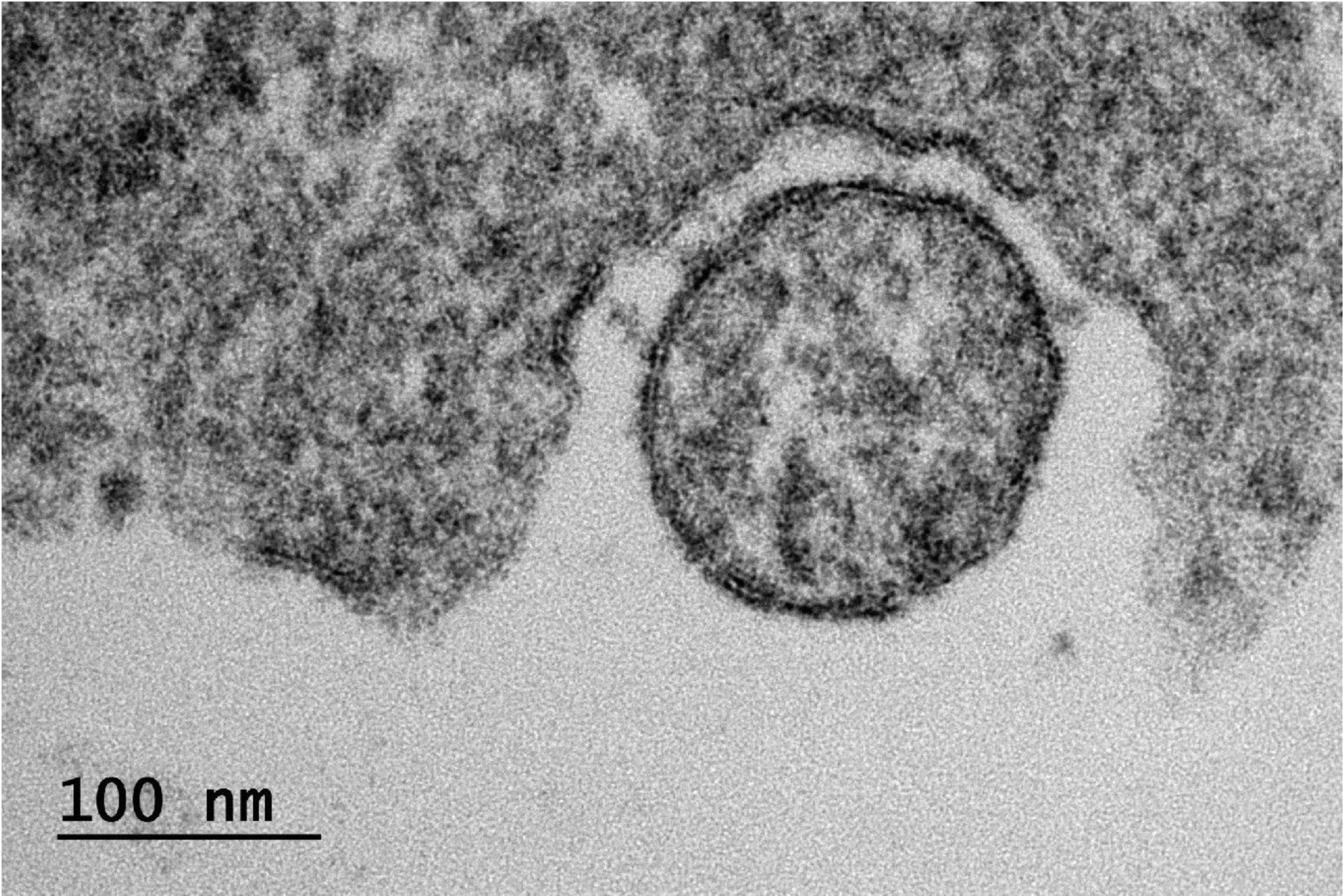
Internalization of a SARS-CoV-2Wu:(E,S,M) observed via thin-section electron microscopy of murine lung epithelial (*MLE*) *cell*s. A VLP can be seen bound to the plasma membrane via the SARS-CoV-2 S protein.

**Extended Data Fig.2.**
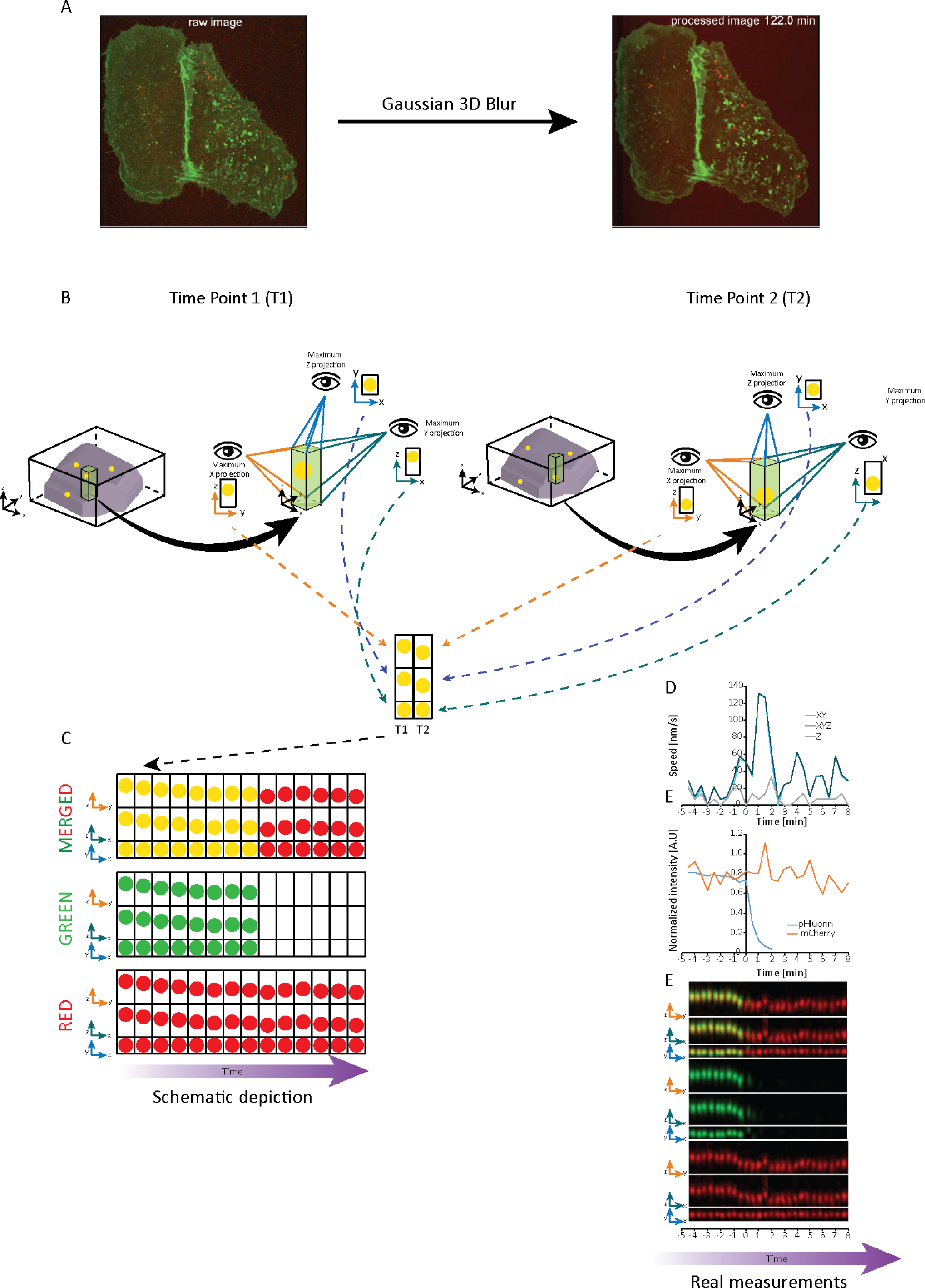
Schematic representation of the SPARTACUSS pipeline for measurement and analysis of VLP internalization dynamics. A. A schematic 3D representation of a cell (gray volume inside black cube) imaged over two consecutive time points. Single VLPs are presented as yellow dots (merged signal mCherry and pHluorin). A volume (green parallelepiped) around a VLP of interest is chosen, and maximum intensity projections along each axis (eye viewpoints) are extracted and combined vertically. The combined projections from the two time points are then horizontally compiled in a sequence to generate a kymograph. B. Schematic kymographs of a single VLP tracked over time, acquired in two imaging channels: green (middle), red (bottom), and merged (top). If the signal in the green channel disappears, the VLP in the merged kymograph changes from yellow to red. For each channel, the different axis projections are presented (ZY - top, ZX-middle, YX - bottom) as kymographs. C. Measurements of the speed of a single VLP using the pipeline described above. Each point in the top graph represents the speed of the particle over time extracted from two consecutive images in 1D-vertically (Z), 2D (XY), and 3D (XYZ) D. Measurement of the intensity of a single VLP for each time point in the two different channels. E. Kymograph of a single SARS-CoV-2 VLP acquired in two channels (Green-M-pHluorin, Red-M-mCherry), presented as described in (B).

**Extended Data Fig.3.**
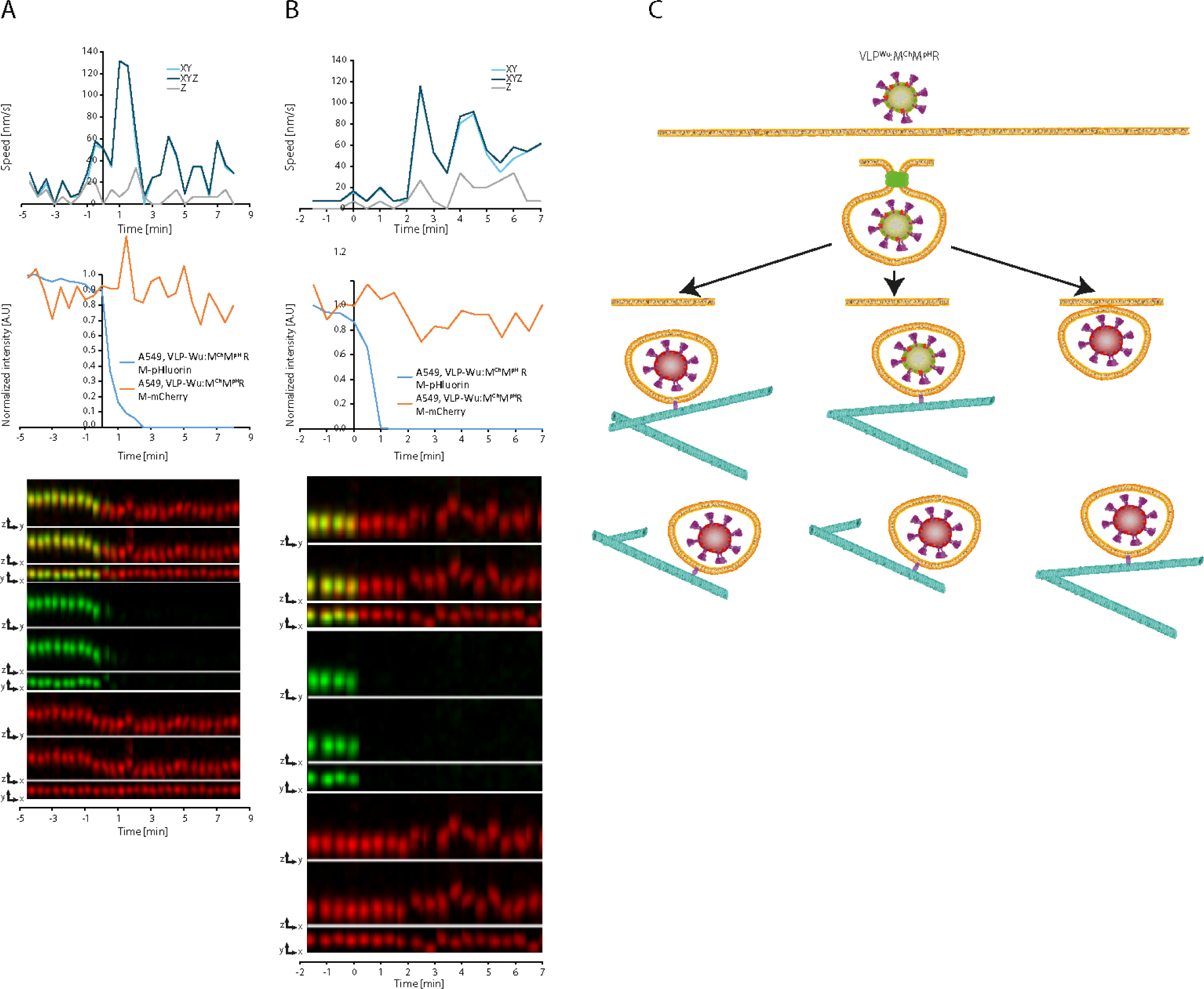
Example of single VLP^Wu^:M^Ch^M^pH^R entries into А549 cells. A. Representative VLP speed and intensity graphs (top) and kymographs (merged, M-mCherry, and pHluorin) in all dimensions (bottom) for a single VLP undergoing internalization, where the speed of the particle increases before its pHluorin intensity starts to decrease. B. Same as (A), but for a VLP the speed of which increases after its pHluorin intensity starts to decrease. C. Schematic representation of the VLP acidification and microtubule attachment (speed increase); The pH drops simultaneously with the start of active microtubule movement (Left); The pH drops below 5 after the start of active microtubule movement (Middle); The pH of the VLP drops below 5 before active microtubule movement (Right);.

**Extended Data Fig.4.**
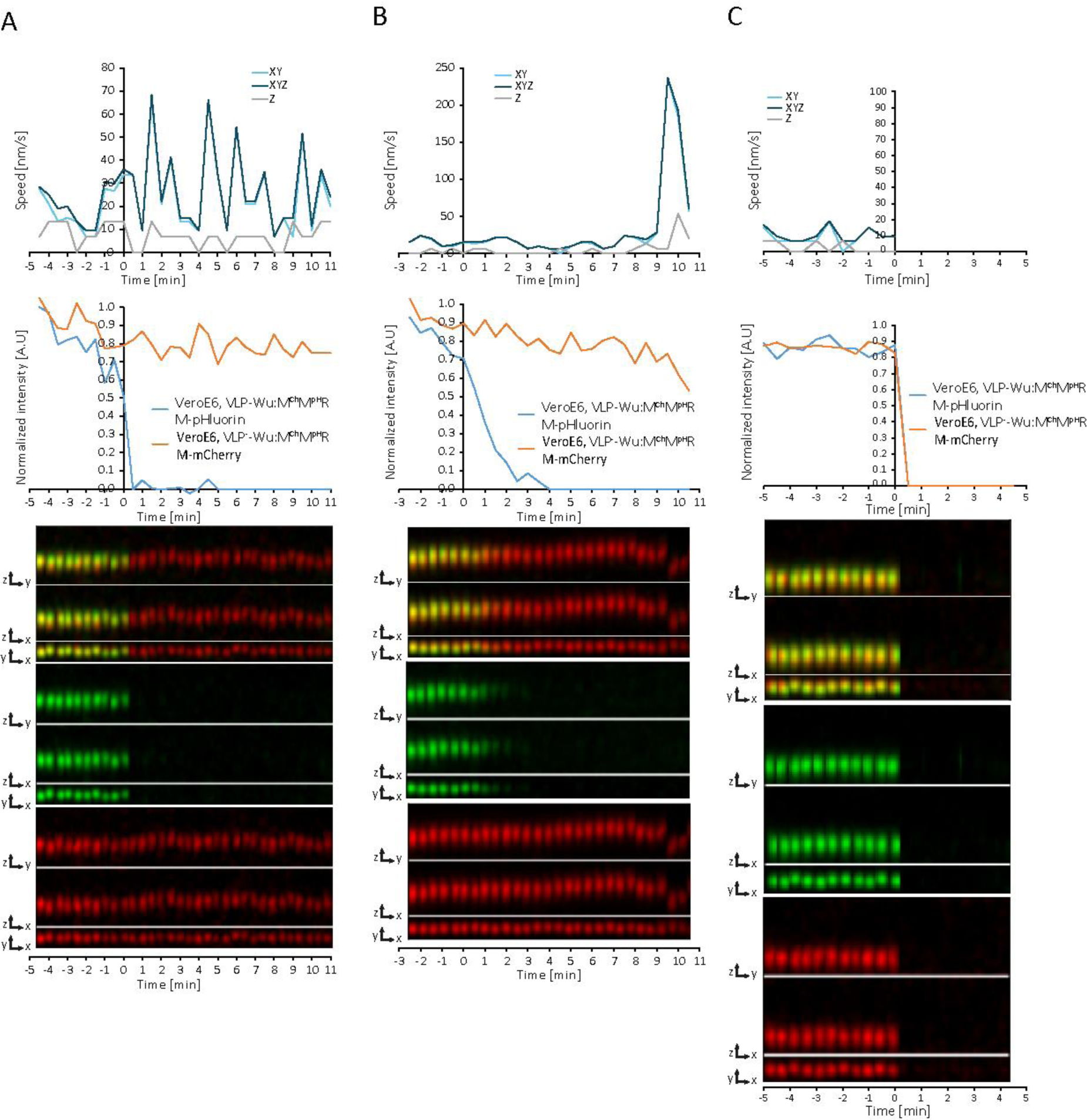
Example of single VLP^Wu^:M^Ch^M^pH^R entries into Vero E6 cells. A. Representative VLP speed and intensity graphs (top) and kymographs (merged, M-mCherry, and pHluorin) in all dimensions (bottom) for a single VLP undergoing internalization, where the speed of the particle increases before its pHluorin intensity starts to decrease. B. Same as (A), but for a VLP the speed of which increases after its pHluorin intensity starts to decrease. C. Same as (A), but for a VLP for which the signals of both fluorescent proteins disappear simultaneously.

**Extended Data Fig.5.**
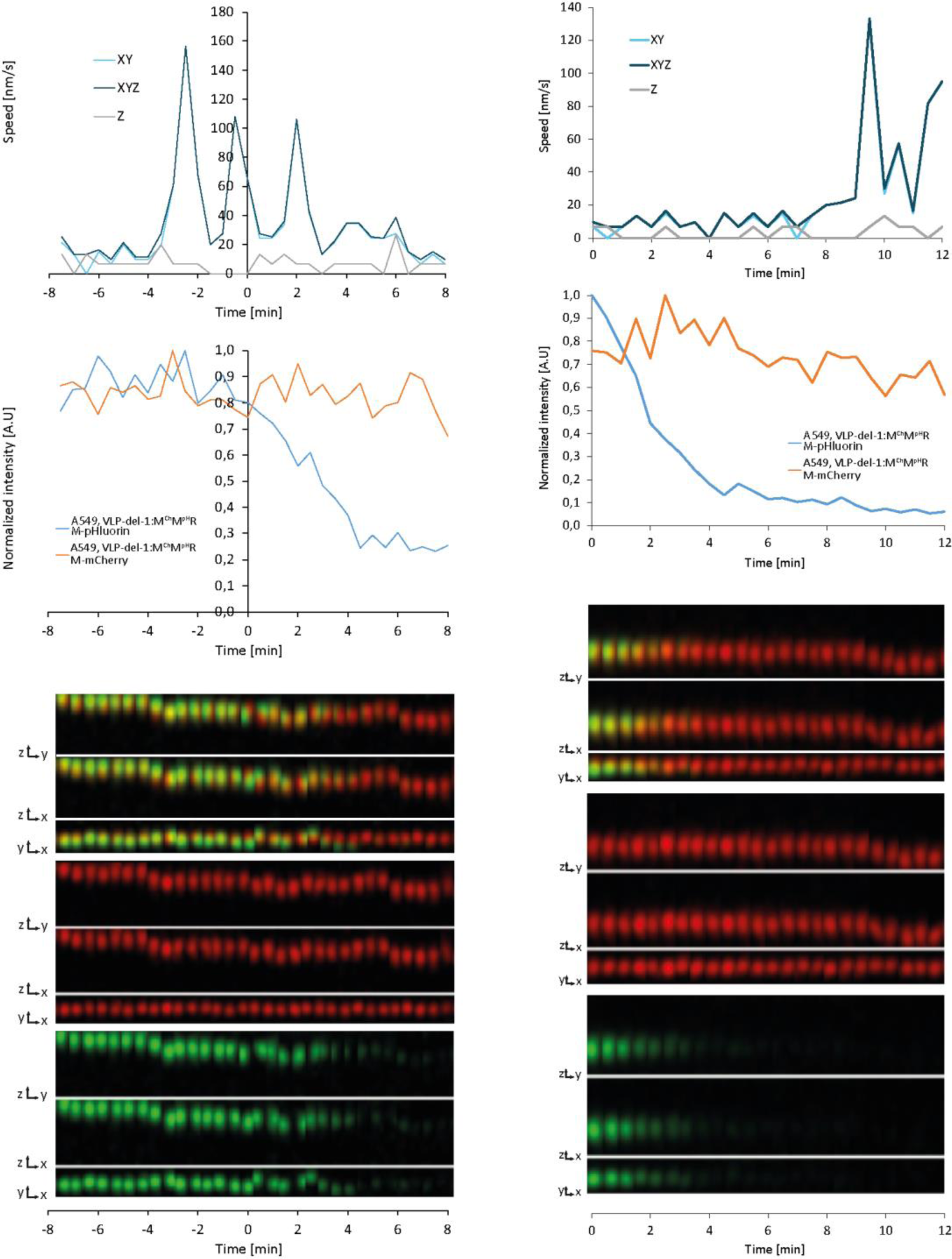
Example of single VLP^del-1^:M^Ch^M^pH^R entry into A549 cells. A. Representative VLP speed and intensity graphs (top) and kymographs (merged, M-mCherry, and pHluorin) in all dimensions (bottom) for a single VLP undergoing internalization, where the speed of the particle increases before its pHluorin intensity starts to decrease. B. Same as (A), but for a VLP the speed of which increases after its pHluorin intensity starts to decrease.

**Extended Data Fig.6.**
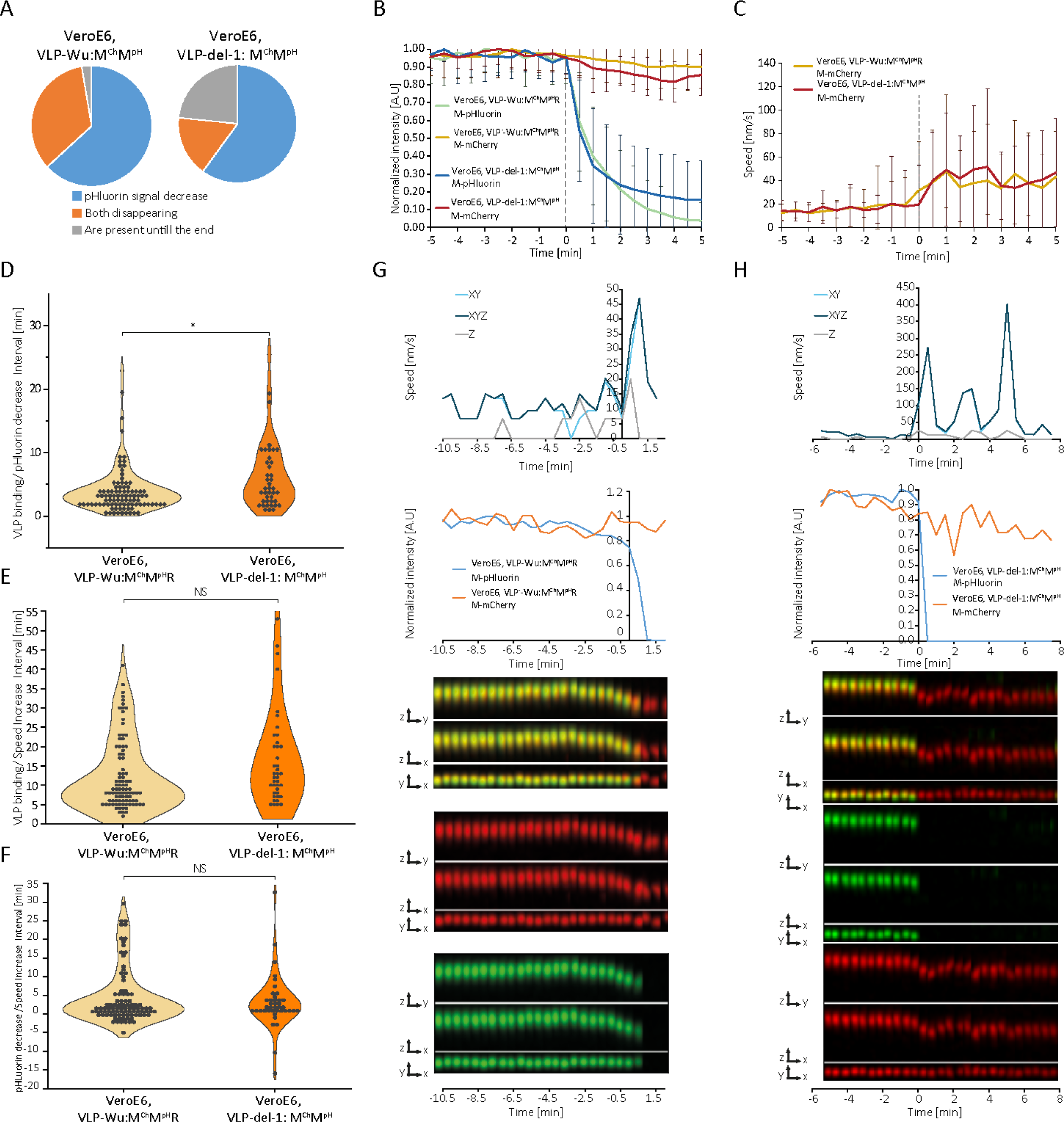
Dynamics of VLP binding and speed increase for VLP^Wu^:M^Ch^M^pH^ and VLP^del-1^:M^Ch^M^pH^ during internalization in VeroE6 cells. A. Percentage of VLPs in which only the M-pHluorin intensity decreases (Blue), M-pHluorin and M-mCherry intensities decrease simultaneously (Orange), or neither decreases (Gray). B. Comparison of pHluorin intensity decrease during the internalization of VLP^Wu^: M^Ch^M^pH^R and VLP^del-1^: M^Ch^M^pH^R in VeroE6 cells. The average intensity of pHluorin is represented as a function of time where the individual VLPs were aligned to the start of VLP pHluorin decrease (0 min). The average M-mCherry intensity is also presented. Error bars represent the standard deviation. For VeroE6-VLP^Wu^:M^Ch^M^pH^R n=93, for VeroE6-VLP^del-1^:M^Ch^M^pH^ n=41. C. Comparison of the dynamics of speed increase of VLP^Wu^:M^Ch^M^pH^ and VLP^del-1^:M^Ch^M^pH^ internalization in Vero E6 cells. The average speed of VLPs is presented as a function of time where the individual VLPs were aligned to the start of M-pHluorin decrease (0 min). Error bars represent the standard deviation. For VeroE6-VLP^Wu^:M^Ch^M^pH^ n=93, for VeroE6-VLP^del-1^:M^Ch^M^pH^ n=41. D. Distribution of time intervals between VLP^Wu^:M^Ch^M^pH^ and VLP^del-1^:M^Ch^M^pH^ binding and start of the decrease in pHluorin intensity during internalization in VeroE6 cells. Two-tailed Student’s t-test; NS p>0.01; * p<0.01. For VeroE6-VLP^Wu^:M^Ch^M^pH^ n=93, for VeroE6-VLP^del-1^:M^Ch^M^pH^ n=41. E. Distribution of time intervals between VLP^Wu^:M^Ch^M^pH^ and VLP^del-1^:M^Ch^M^pH^ binding and start of VLP speed increase in VeroE6 cells. Two-tailed Student’s t-test; NS p>0.01; * p<0.01. For VeroE6-VLP^Wu^:M^Ch^M^pH^ n=93, for VeroE6-VLP^del-1^:M^Ch^M^pH^ n=41. F. Distribution of time intervals between VLP^Wu^:M^Ch^M^pH^ and VLP^del-1^:M^Ch^M^pH^ pHluorin intensity decrease and start of VLP speed increase in VeroE6 cells. Two-tailed Student’s t-test; NS p>0.01; * p<0.01. For VeroE6-VLP^Wu^:M^Ch^M^pH^ n=93, for VeroE6-VLP^del-1^:M^Ch^M^pH^ n=41. G. Representative VLP speed and intensity graphs (top) and kymographs (merged, M-mCherry, and pHluorin) in all dimensions (bottom) for a single VLP^Wu^:M^Ch^M^pH^ undergoing internalization in a VeroE6 cell. H. Same as (G) for VLP^del-1^:M^Ch^M^pH^.

**Extended Data Fig. 7.**
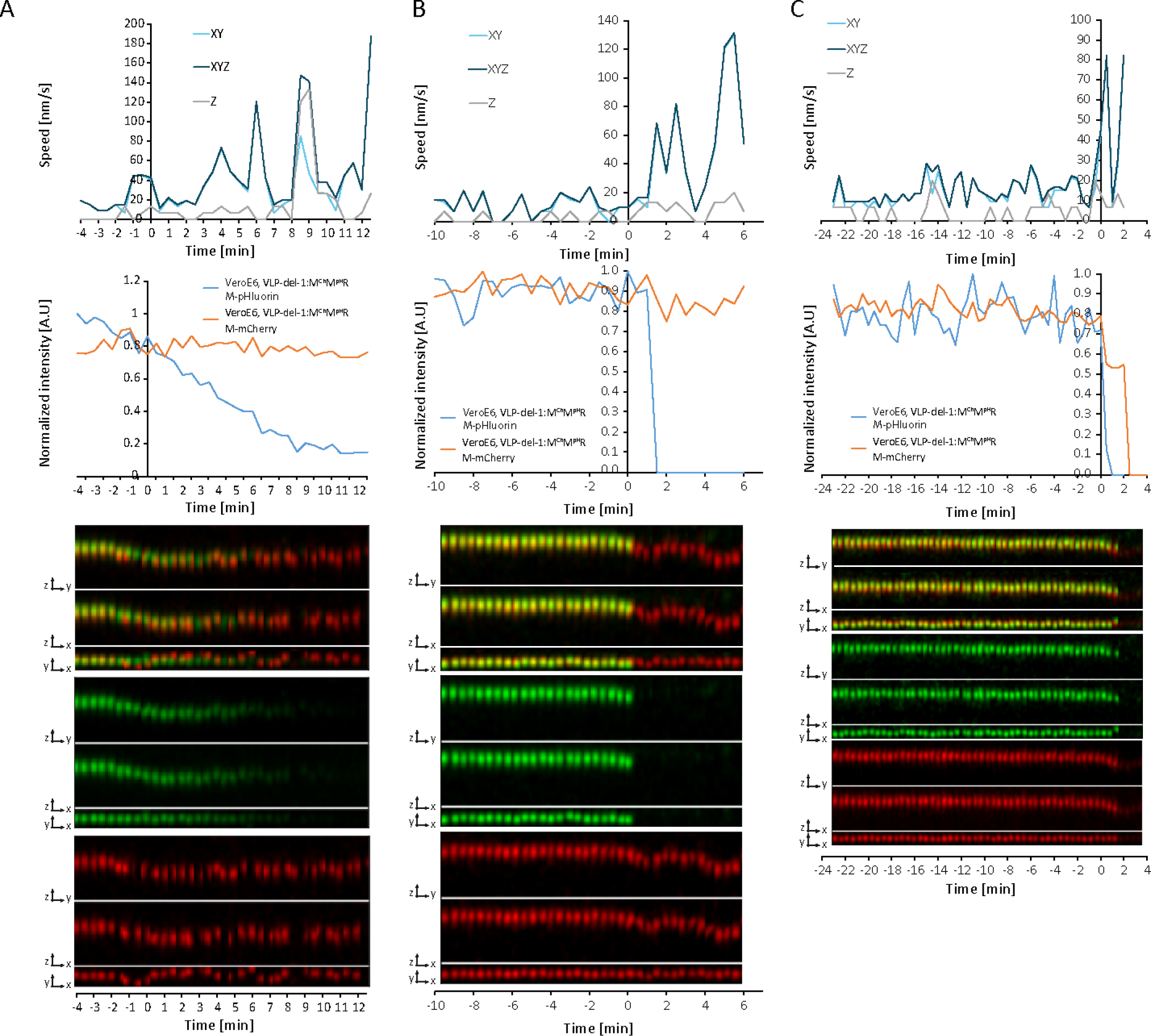
Examples of a single VLP^del-1^:M^Ch^M^pH^ entry into Vero E6 cells. A. Representative VLP speed and intensity graphs (top) and kymographs (merged, M-mCherry, and pHluorin) in all dimensions (bottom) for a single VLP undergoing internalization, where the speed of the particle increases before its pHluorin intensity starts to decrease. B. Same as (A), but for a VLP the speed of which increases after its pHluorin intensity starts to decrease. C. Same as (A), but for a VLP for which the signals of both fluorescent proteins disappear simultaneously.

**Extended Data Fig. 8.**
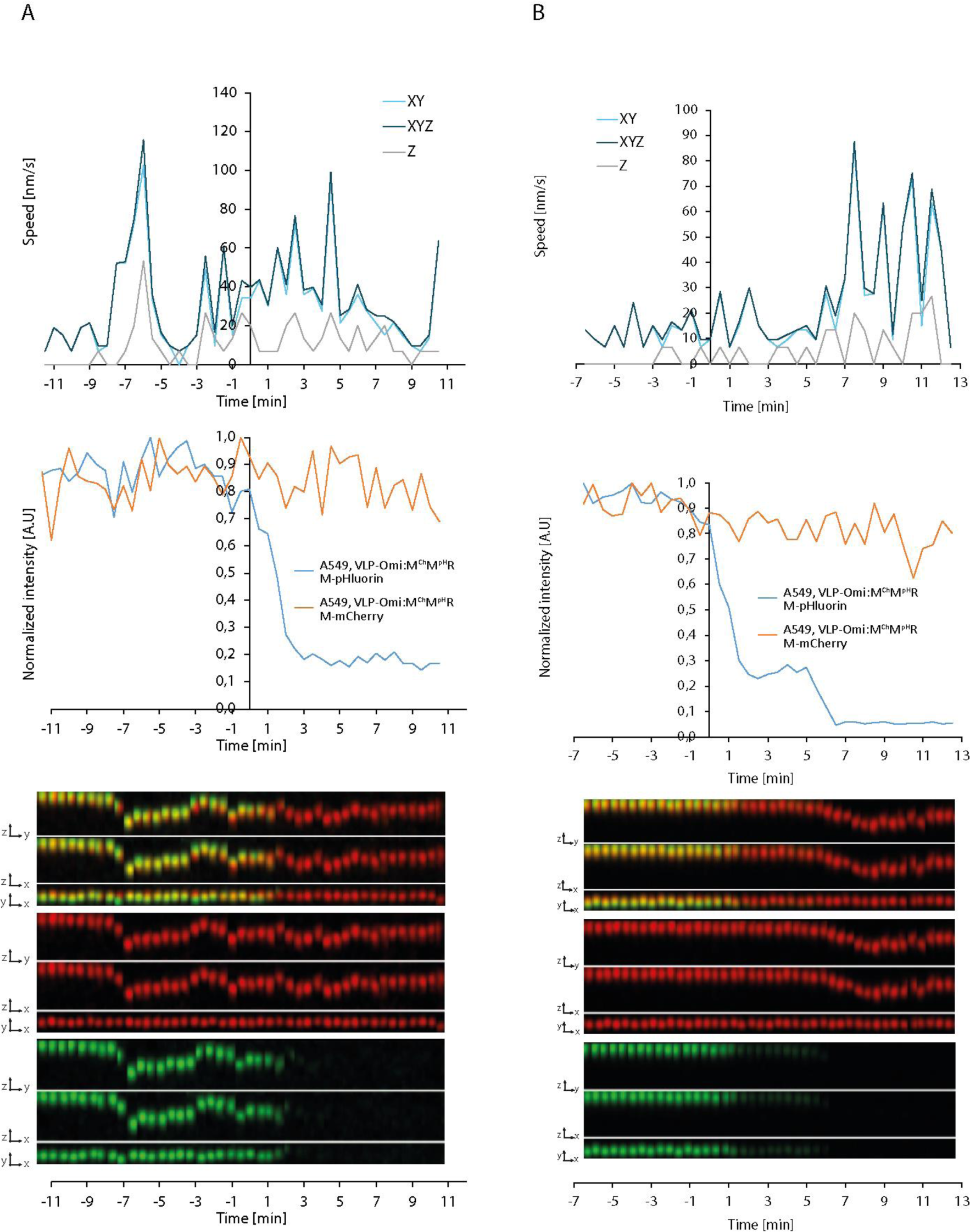
Examples of single VLP^Omi^:M^Ch^M^pH^R entries into A549 cells. A. Representative VLP speed and intensity graphs (top) and kymographs (merged, M-mCherry, and pHluorin) in all dimensions (bottom) for a single VLP undergoing internalization, where the speed of the particle increases before its pHluorin intensity starts to decrease. B. Same as (A), but for a VLP the speed of which increases after its pHluorin intensity starts to decrease.

**Extended Data Fig.9.**
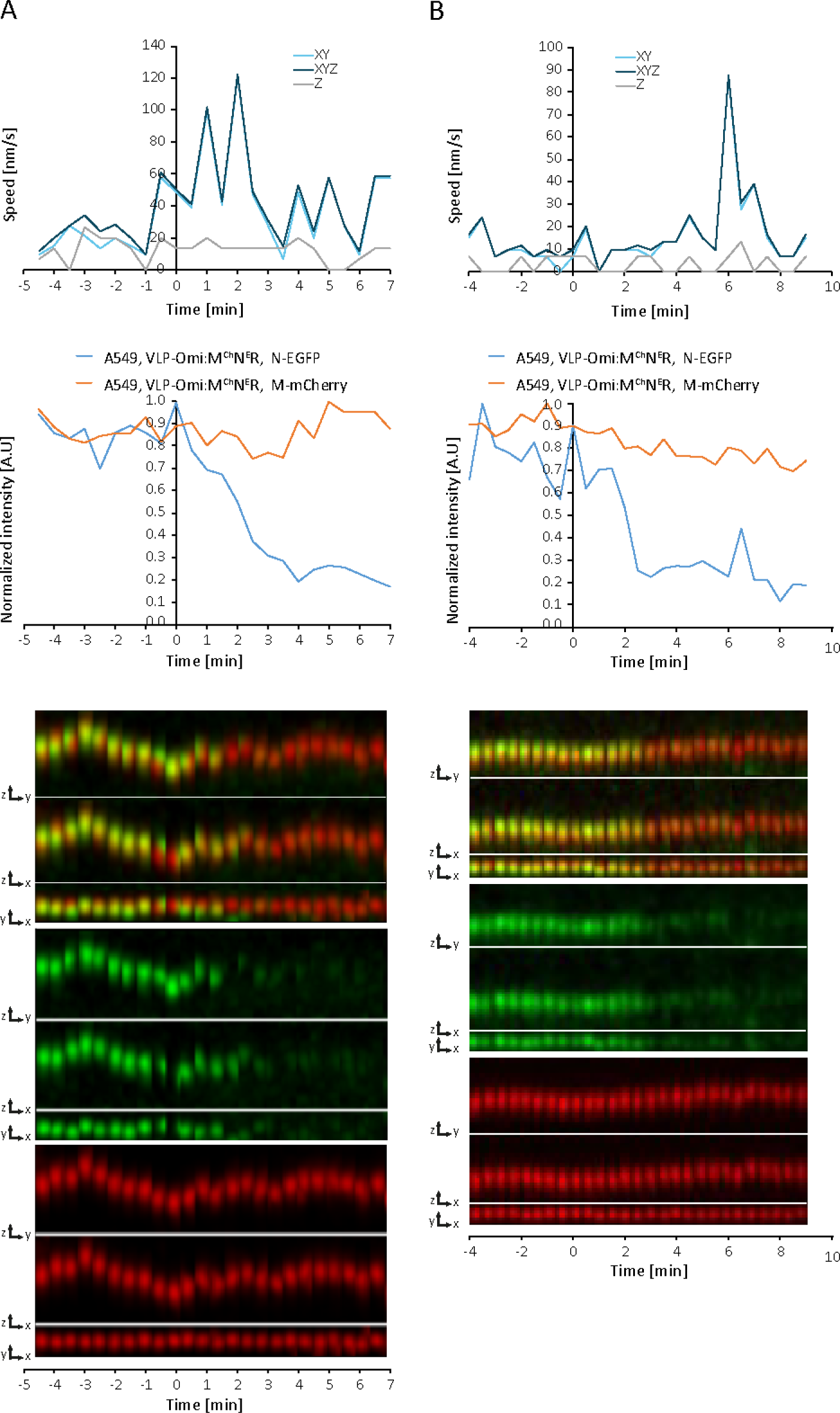
Examples of single VLP^Omi^:M^Ch^N^E^R entries into А549 cells. A. Representative VLP speed and intensity graphs (top) and kymographs (merged, M-mCherry, and N-EGFP) in all dimensions (bottom) for a single VLP undergoing internalization, where the speed of the particle increases before its pHluorin intensity starts to decrease. B. Same as (A), but for a VLP the speed of which increases after its pHluorin intensity starts to decrease.

**Extended Data Fig. 10.**
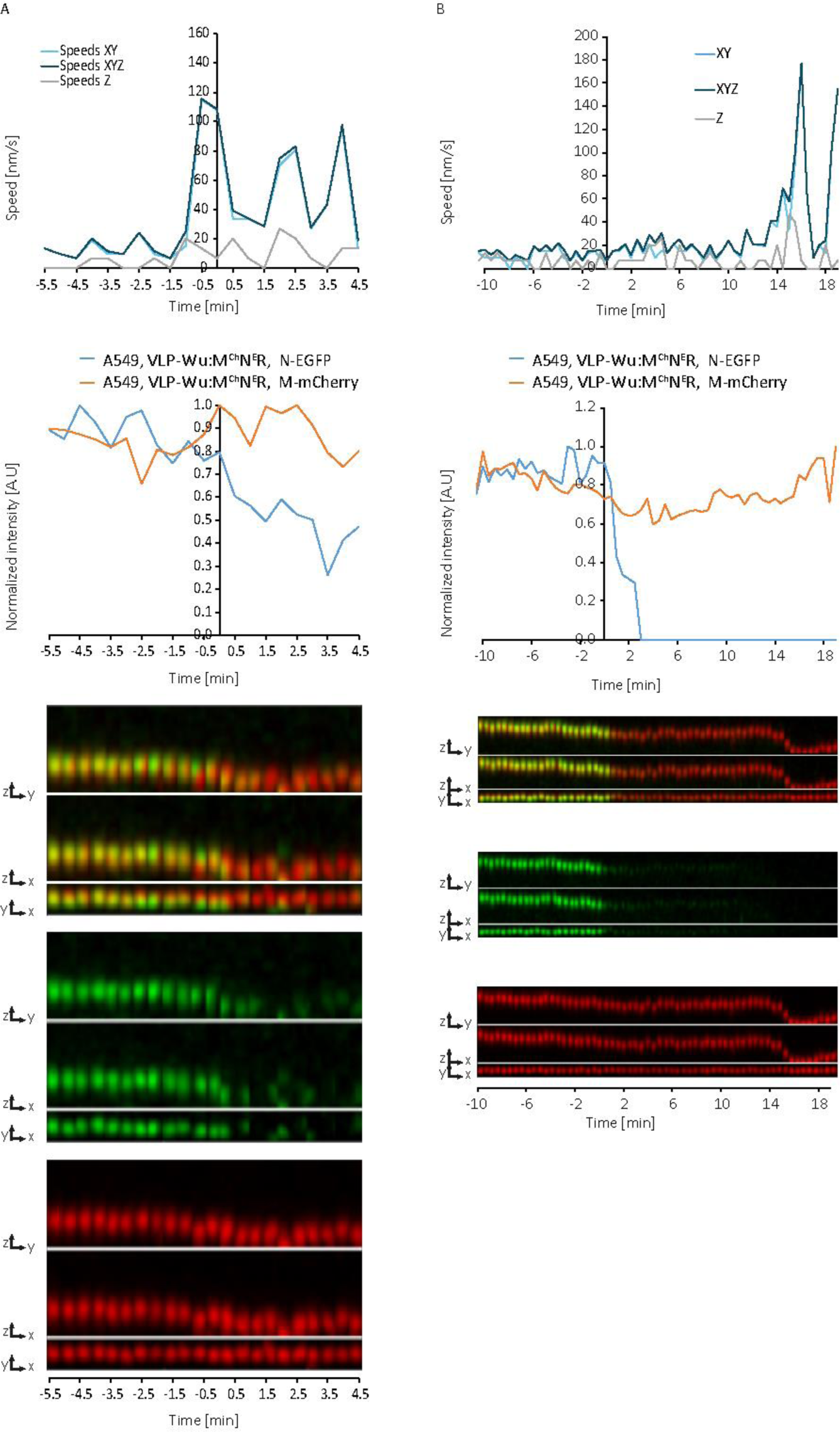
Examples of single VLP^Wu^:M^Ch^N^E^R entries into А549 cells. A. Representative VLP speed and intensity graphs (top) and kymographs (merged, M-mCherry, and N-EGFP) in all dimensions (bottom) for a single VLP undergoing internalization, where the speed of the particle increases before its pHluorin intensity starts to decrease. B. Same as (A), but for a VLP the speed of which increases after its pHluorin intensity starts to decrease.

